# Why amyloid fibrils have a limited width

**DOI:** 10.1101/2021.07.02.450971

**Authors:** David R. Boyer, Nikos A. Mynhier, Michael R. Sawaya

**Affiliations:** Department of Chemistry and Biochemistry and Biological Chemistry, UCLA-DOE Institute, Molecular Biology Institute and Howard Hughes Medical Institute, UCLA, Los Angeles, CA, USA; Department of Cell Biology, Blavatnik Institute, Harvard Medical School, Boston, MA, USA; Department of Pediatric Oncology, Dana-Farber Cancer Institute, Boston, MA, USA

**Keywords:** Amyloid fibril, cryo-EM, fibril structure, amyloid width, cross-β, amyloid hydrogen bonding

## Abstract

Amyloid fibrils can grow indefinitely long by adding protein chains to the tips of the fibril through β-sheet hydrogen bonding; however, they do not grow laterally beyond ∼10-20 nm. This prevents amyloid fibrils from growing into two-dimensional or three-dimensional arrays. The forces that restrict lateral association of β-sheets in amyloid fibrils are not immediately apparent. We hypothesize that it is the helical symmetry of amyloid fibrils that imposes the limit on fibril width by incurring an increasing separation between helically related molecules as a function of radial distance from the helical axis. The unavoidable consequence is that backbone hydrogen bonds that connect symmetrically related layers of the fibril become weaker towards the edge of the fibril, ultimately becoming too weak to remain ordered. To test our hypothesis, we examined 57 available cryo-EM amyloid fibril structures for trends in interstrand distance and β-sheet hydrogen bonding as a function of radial distance from the helical axis. We find that all fibril structures display an increase in interstrand distance as a function of radius and that most fibril structures have a discernible increase in β-sheet hydrogen bond distances as a function of radius. In addition, we identify a high resolution cryo-EM structure that does not follow our predicted hydrogen bonding trends and perform real space refinement with hydrogen bond distance and angle restraints to restore predicted hydrogen bond trends. This highlights the potential to use our analysis to ensure realistic hydrogen bonding in amyloid fibrils when atomic resolution cryo-EM maps are not available.

**Significance Statement:** The number of amyloid fibril structures determined has exploded in recent years due to advances in structural biology techniques. However, we are still at the beginning stages of understanding amyloid fibril assembly. One important property that is critical to fibril formation and mechanical properties is the fibril width. Despite the diversity of fibril folds discovered, all amyloid fibrils are constrained to a width of 10-20 nm. Here, we use simple geometry and structural analysis to identify that the limited width of amyloid fibrils arises from the helical twist of β-sheets in amyloid fibrils. Our findings provide important considerations for the accurate modeling of hydrogen bonds in amyloid fibrils as well as for the possible prediction and design of amyloid-based nanomaterials.

## Introduction

A wide variety of protein sequences can form amyloid fibrils, including proteins that fibrillize as part of their biological function or in disease, as well proteins that only fibrillize under non-biological conditions in the test tube^1, 2^. However, even though the sequences and structural details of individual amyloid fibrils can be quite different, all amyloid fibrils share a common blue-print^3^. To a first approximation, amyloid fibrils are composed of a single protein sequence that adopts a largely two-dimensional, serpentine-like fold. This fold has two exposed surfaces – the top and bottom of the two-dimensional layer – that stack repeatedly with other identical two-dimensional layers with a slight helical twist. The fibril can extend indefinitely along the helical axis by adding further self-interacting protein chains to the “open” top and bottom layers of the fibril. This open-ended assembly differs from most protein assemblies, which are “closed”, globular, and discrete entities with a finite, invariable number of subunits.

The architectural features of amyloid fibrils described above, as well as further details about the fibril structure, are encapsulated in the term cross-β fold. The cross-β fold describes not only the interactions that stabilize the subunits in the fibril (quaternary structure), but also the interactions between different sections of the same protein chain (tertiary structure) and the conformation of contiguous sections of the protein chain (secondary structure). In the cross-β secondary structure, the amino acids in amyloid fibrils largely adopt β-strand conformations where the backbone carbonyl oxygens and amide hydrogens point alternately up and down the fibril axis. In the tertiary structure, β-strands are punctuated by turns that allow the sequence to fold back on itself and form mated β-strands whose side chains interact in the direction orthogonal to the fibril axis. Finally, the cross-β quaternary structure is formed by individual two-dimensional protein chains stacking upon each other along the fibril axis, held together by the hydrogen bonding of the main chain carbonyl oxygens and amide hydrogens, into parallel, in-register β-sheets.

Given that lateral interactions between mated β-sheets in the amyloid fibril are pervasive among amyloid fibrils, it is a mystery why amyloid fibrils do not grow wider than a certain amount. Indeed, in crystal structures of amyloid peptides, we often observe that the cross-β structure can extend many thousands of copies in the plane orthogonal to the fibril axis^4^. This is achieved through repeated lateral mating of β-sheets via their side chains in much the same way the β-sheets grow indefinitely along the fibril axis through repeated mainchain hydrogen bonding in both crystals and fibrils. Figure 1 illustrates the differences between crystals and fibrils by comparing the crystal structure of the longest amyloid peptide crystallized to date – Aβ 20-34 isoasp23^5^ – with an amyloid fibril structure of a full-length amyloid protein – Aβ 1-40^6^. In the crystal lattice, the addition of peptides occurs indefinitely through side chain interactions in the entire plane orthogonal to the fibril axis, while in the fibril structure, the sides of the fibril structure are exposed and available for additional lateral interactions, yet no additional protein adds to the fibril laterally. In both cases, additional layers add along the fibril axis (extending in and out of the page) indefinitely through mainchain hydrogen bonding of β-sheets.

**Figure 1.**
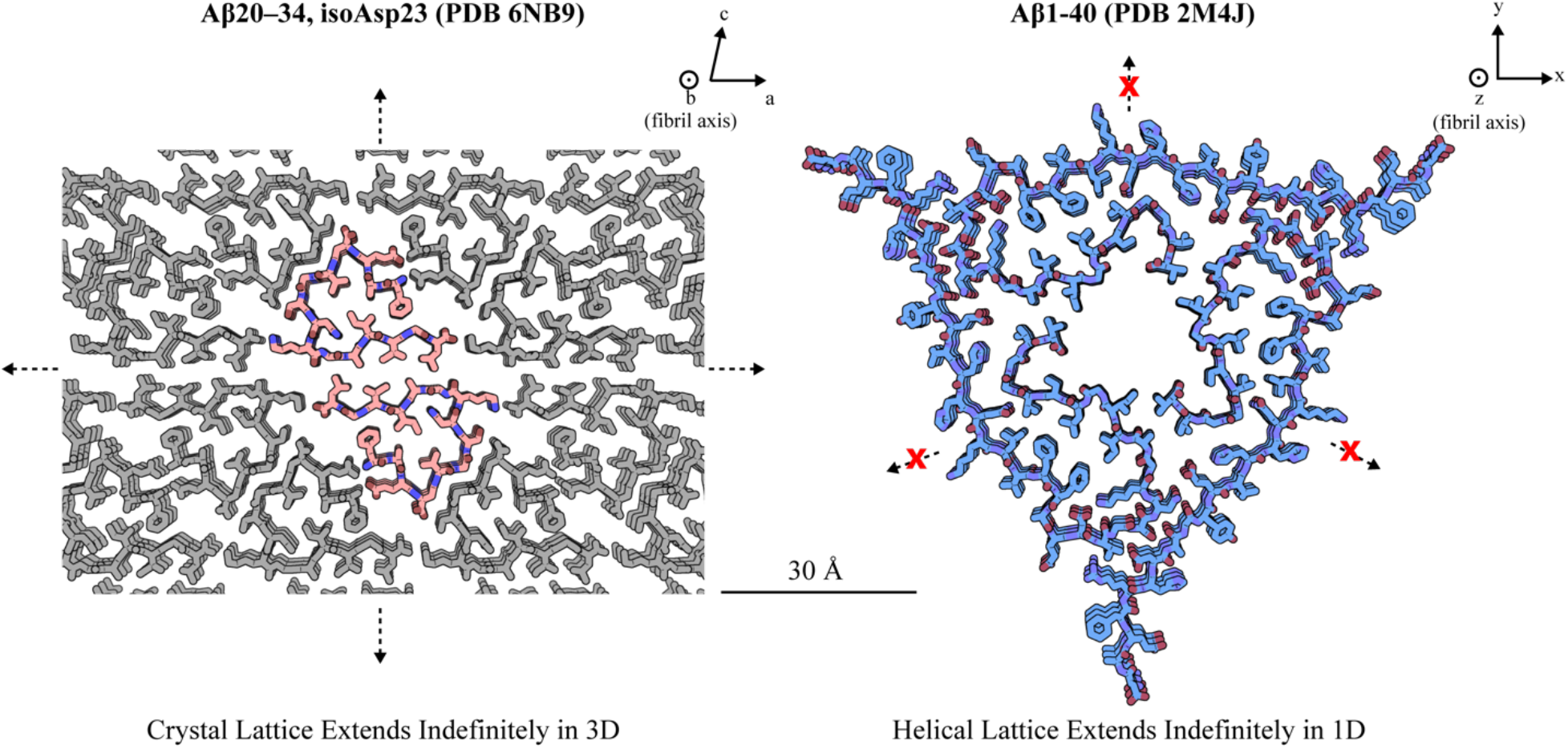
Comparison between Aβ(20-34) isoasp23 crystal structure and Aβ(1-40) fibril structure. Peptide molecules in the crystal lattice repeat indefinitely in all dimensions due to lateral side chain-side chain interactions as well as backbone hydrogen bonding along the fibril axis. The fibril does not grow laterally through repeated mating of additional protofilaments, and instead, extends indefinitely along the fibril axis through backbone hydrogen bonding, similar to the crystal lattice.

There are two possible mechanisms for amyloid fibrils to grow wider: i) additional protofilaments can be added to exposed surfaces of the fibril, ii) additional residues of the protein chain add to the cross-β fold. The former case is akin to that of a short peptide crystal structures where repeated addition of protein chains occurs orthogonal to the fibril axis, while the latter case is only applicable to fibrils since in amyloid fibrils formed from full-length protein there are generally only a certain subset of amino acids in the protein chain that participate in forming the cross-β fold of the fibrils – the “fibril core”. Residues not in the fibril core form the “fuzzy coat” – a disordered tangle of amino acids coating the length of the fibril. The fact that additional protofilaments do not add indefinitely to the fibril and that many residues are found in the fuzzy coat add to the mystery of why amyloid fibrils do not grow wider.

A key feature of amyloid fibrils is their helicity, which arises from the twisting of the β-strands in each protein chain. Twisted β-strands generate twisted β-sheets, due to the fact that the β-strands in a β-sheet have a repeated, asymmetric interface^7^. In general, a right-handed β-strand twist is thought to be more energetically stable for L-amino acids, and this gives rise to a left-handed β-sheet twist^8^. However, there are also examples of left-handed β-strands and corresponding right-handed β-sheets in nature^9^. Figure 2 shows a prototypical left-handed double helix where the relationship between symmetrically related subunits is given by two parameters: the helical *twist*, in °/subunit, which describes the incremental rotation of each subunit around the helical axis and the helical *rise*, in Å/subunit, which describes the translation of the subunits along the helical axis. In cylindrical coordinates, the relationship between identical objects in a helix is given by:

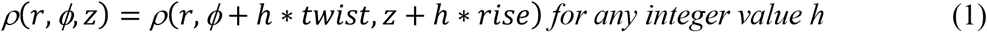

**Figure 2.**
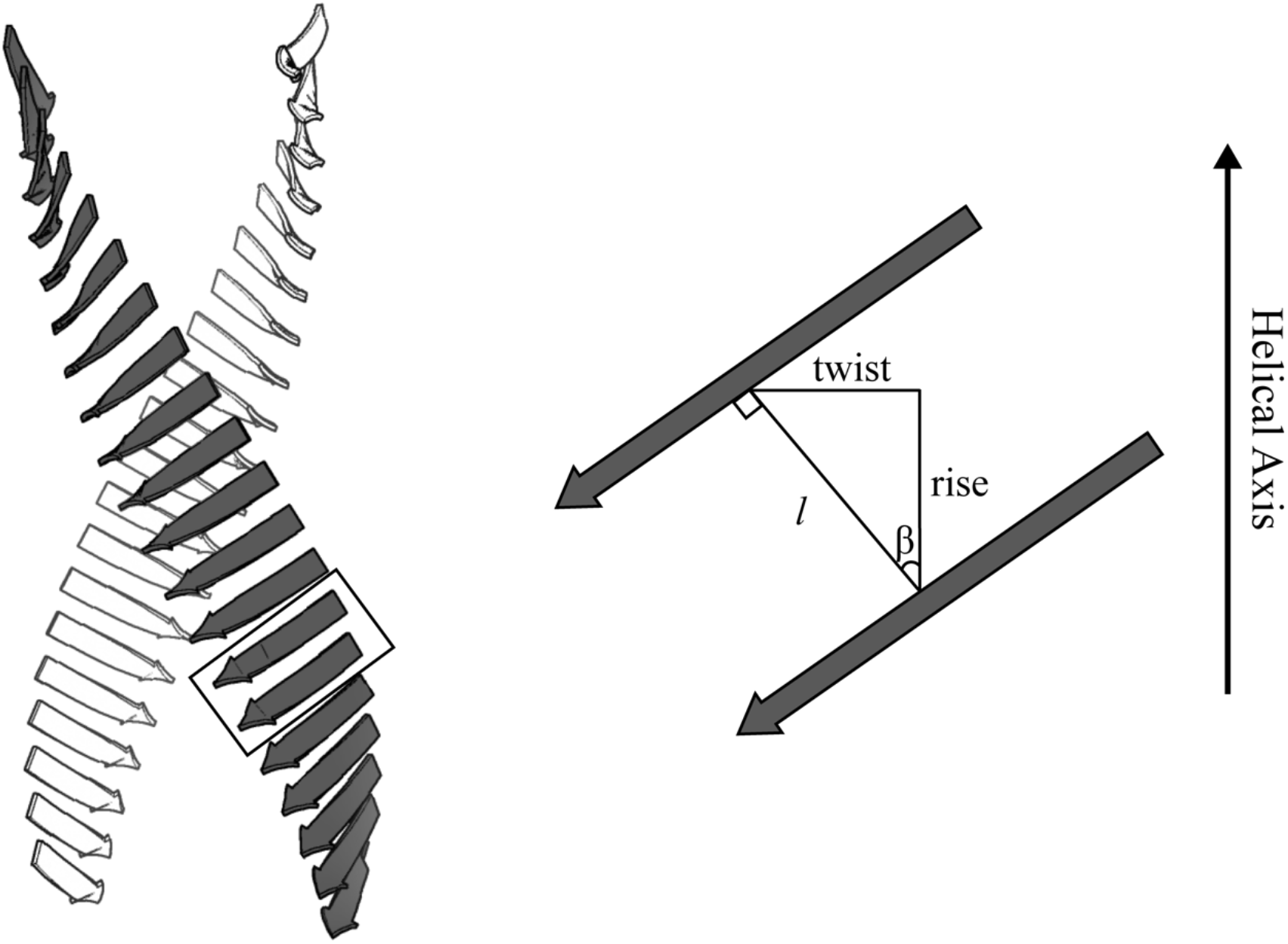
The relationship between subunits in a helix can be described by an azimuthal twist around the helical axis and a translational rise along the helical axis. The displacement between adjacent subunits is *l*, which can be calculated with Eq. 3. In order to maintain hydrogen bond geometry between adjacent β-strands in an amyloid fibril, the strands tilt according to Eq. 4.

Where *r* is the radial distance from the helical axis, *ϕ* is the angular coordinate, and *z* is the height. And the distance between two objects in cylindrical coordinates is:

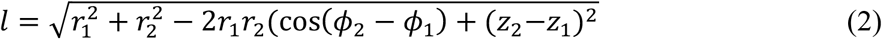

Which reduces to Eq. 3 when calculating the distance between subunits i and i+1 (two consecutive layers in a helix, Figure 2):

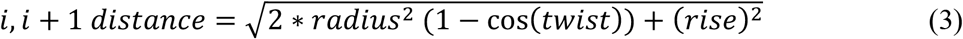

Eq. 3 shows that the distance between the *i* and *i+*1 subunits is proportional to their radial distance from the helical axis (*radius*). In addition, Eq. 3 demonstrates that the *i* to *i+*1 distance is dependent on the twist and rise parameters of the helix. In the case of amyloid fibrils, the rise is invariably ∼4.8 Å per layer; however, the twist can vary for each fibril. Fig. 3 shows the relationship between the i to i*+*1 distance and radius for different twist values. The *i* to *i*+1 distance increases with longer radius according to Eq. 3, and the rate of increase is steeper at higher twist values.

**Figure 3.**
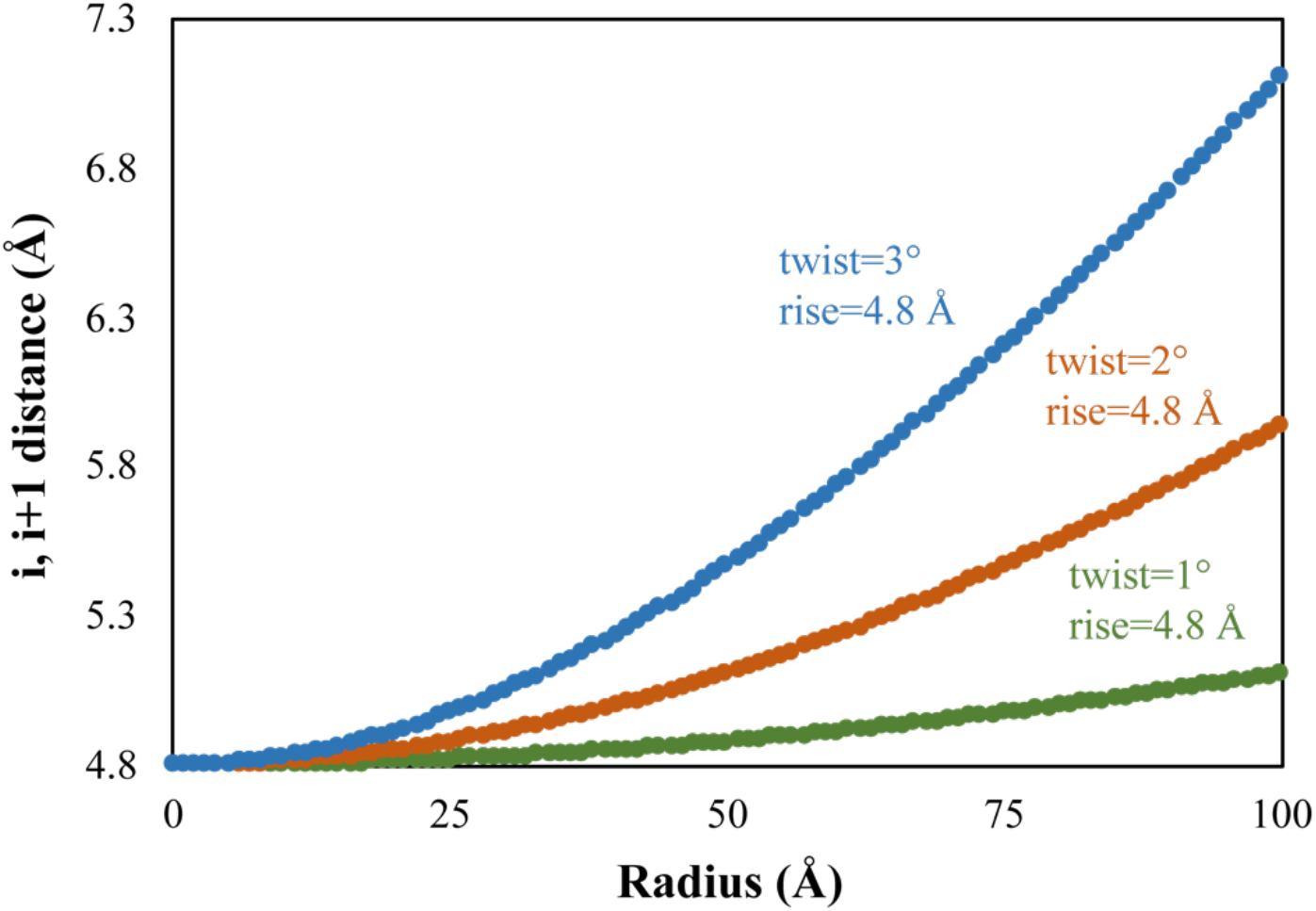
Simulated *i* to *i*+1 distances versus radial distance from the helical axis (radius) for different twist values. For helices with a larger twist, the distance between adjacent subunits increases more quickly as a function of radial distance from the helical axis.

We hypothesize that the helical symmetry of amyloid fibrils places a restraint on the maximum width of the fibril. As the distance between the i to i+1 β-strands increases towards the edge of the fibril, they will not be close enough for β-sheet hydrogen bonding and will not be able to form part of the ordered core. Furthermore, as Eq. 3 and Figure 3 show, fibrils with a larger twist will have a smaller maximum width since the i to i+1 distance will increase more rapidly. Conversely, fibrils with a smaller twist can grow to a larger maximum width.

Another consequence of the helical symmetry of amyloid fibrils is that as the β-strands get farther from the helical axis, they will have to tilt about the axis orthogonal to the fibril axis in order to maintain the inter-strand hydrogen bond geometry. In Figure 2, this tilt is depicted as the β angle between the fibril axis and the line connecting identical parts of the subunits in the helix. As the radial distance from the helical axis grows, and the distance *l* between i to i+1 subunits grows, so will β according to Eq. 4:

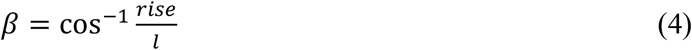

In order to illustrate how the helicity of amyloid fibrils affects the orientation of β-sheets as a function of distance from the helical axis, we transformed the crystal structure of tau peptide SVQIVY into a helix (Figure 4) with a 1°/subunit twist (a twist typical of amyloid fibrils). To modify the helix width, we applied the crystallographic symmetry found in the SVQIVY crystal structure^10^ to add consecutive peptides laterally outward from the helical axis to a maximum radial distance of 125 Å, giving the fibril a diameter of 250 Å. Then, we applied helical symmetry to one layer of the fibril to generate the full helix. We made two copies of the helix: one where the strands in the starting layer were left un-tilted and one where the strands were tilted according to Eq. 4.

**Figure 4.**
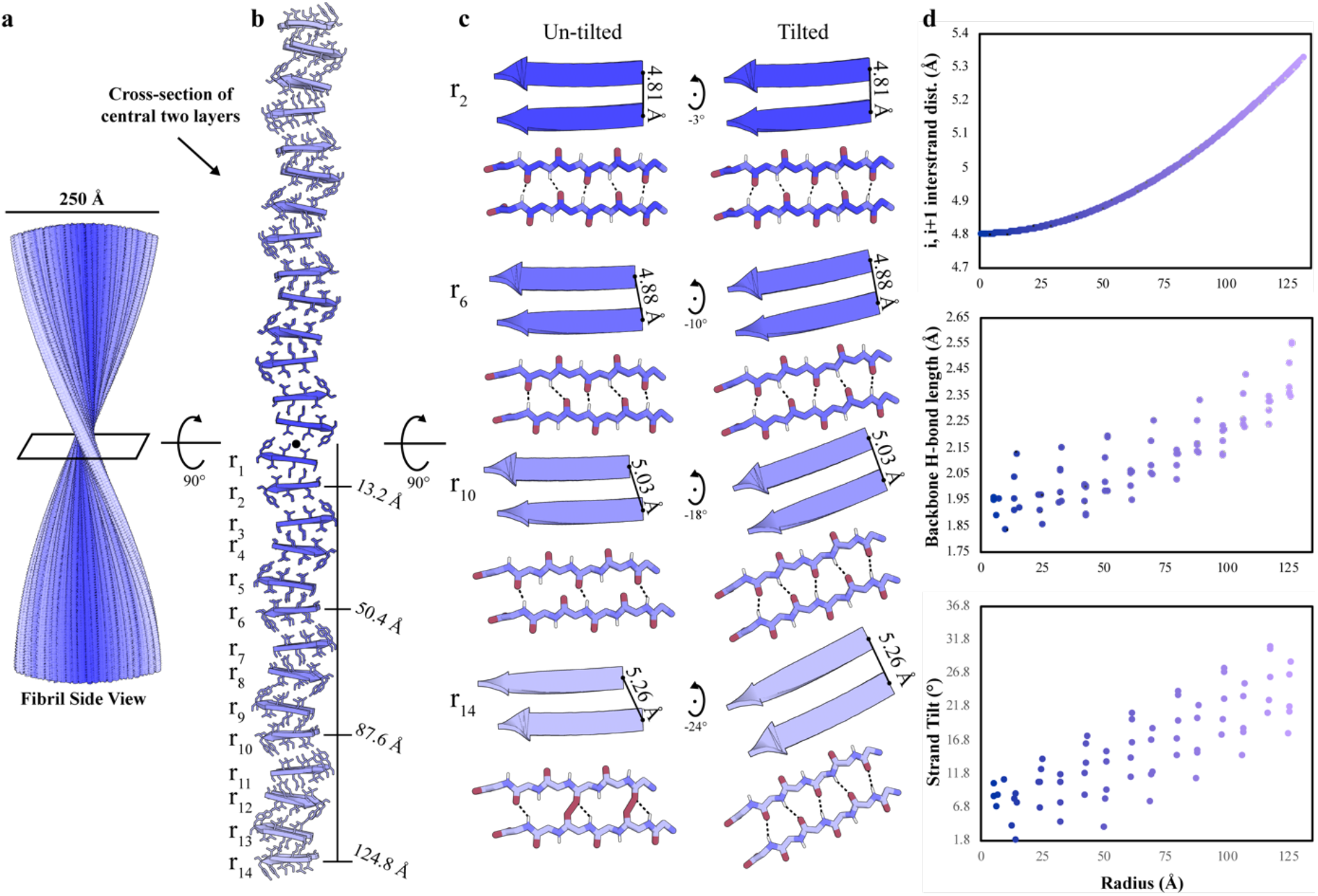
A theoretical, helical amyloid fibril built by adding strands of peptide SVQIVY laterally outward from the fibril axis, built with a 1°/subunit twist and 4.8 Å rise. For all panels, darker color indicates a small distance from the helical axis, while lighter color indicates a large distance from the helical axis. a) Side view of the SVQIVY helix. b) Central cross-section of the helix highlighting the radii of different protofilaments of the fibril. c) Side-view of selected protofilaments at increasing radius from the helical axis – r_2_ (radius = 13.2 Å), r_6_ (radius = 50.4 Å), r_10_ (radius = 87.6 Å), and r_14_ (radius = 124.8 Å) – demonstrates that the distance between strands grows larger as the radius increases and that β-strands need to tilt to maintain hydrogen bond geometry as the radius increases. d) Quantification of symmetry related i to i+1 atoms, β-sheet hydrogen bond lengths and off-axis tilts as a function of radius from helical axis.

Figure 4 shows that as β-sheets lie farther from the helical axis, the distance between the i and i+1 atom increases according to Eq. 3. Also, it is clear that hydrogen bond geometry would not be maintained unless the strands were tilted counterclockwise around an axis orthogonal to the fibril axis (Figure 4c). The top graph in Figure 4d shows the measured i to i+1 distances for all atoms as a function of radial distance from the helical axis while the middle and bottom graphs show the measured hydrogen bond distances (O to H) and β-strand tilts as a function of radial distance from the helical axis, respectively. As expected, the hydrogen bond distances and tilts for the i to i+1 strands increase with increasing interstrand distance as a consequence of helical twist.

The helix of Fig. 4 is only theoretical. In real life, strands would have been unable to associate laterally to the helix at a radius of ∼70 Å, where the i to i+1 distance becomes greater than ∼4.9 Å, the backbone hydrogen bonding in the β-sheets becomes greater than ∼2-2.2 Å, and the strand tilt becomes greater than 15°. Therefore, at least in our thought experiment, the helicity of amyloid fibrils limits their ability to grow wider than a certain amount.

## Results

### Empirical Evidence that Fibril Twist and Radius are Correlated

In order to examine whether the trends we predict are observed in real-life amyloid fibril structures, we conducted an analysis of 57 cryo-EM amyloid fibril structures determined to date (see Supplementary Table 2, Pick’s Disease Wide Fibril and RipK3 used in this analysis, see Methods). We selected cryo-EM structures as opposed to those determined by solid state NMR since the helical twist and rise are known accurately in cryo-EM helical reconstructions. For the 57 structures, we first tested for a correlation between fibril width and fibril pitch. We plotted fibril width against fibril pitch, instead of twist, since pitch takes into account both the helical twist and rise of the structure (for an n-start helix where the entire 4.8 Å layer is considered the helical subunit, pitch=360° * rise / twist). According to Eq. 3 and Figure 3, if fibrils have a large twist (and correspondingly small pitch), the distance between symmetrically related i to i+1 atoms will increase more quickly as the radial distance increases. This should lead to fibrils with a large twist (and small pitch) having a smaller radius. Similarly, if fibrils have a small twist (and correspondingly large pitch), the distance between symmetrically related i to i+1 atoms will increase more slowly as the radial distance increases. This should lead to fibrils with a small twist (and large pitch) having a larger radius.

Figure 5 demonstrates that, in general, wide fibrils will have small twists (and large pitches) and narrow fibrils will have large twists (and small pitches). However, the R^2^ value of 0.5577 demonstrates this is not a perfect correlation. Inspecting the graph, it can be seen that the strongest deviations from the trendline occur when fibrils have a larger pitch (smaller twist) than expected from their radius. This is likely due to the fact that helical symmetry only indicates the upper limit on the helical twist for a fibril of a given radius and does not impose a lower bound on helical twist for a fibril of a given radius. Figure 5 is also consistent with the correlation found by Wu, et al. between the number of residues in the fibril core (a proxy for fibril width) versus fibril pitch when they correctly identified that the very small core of RIPK3 fibrils in their study allowed for a large helical twist (the bottom- and left-most point on Figure 5)^11^.

**Figure 5.**
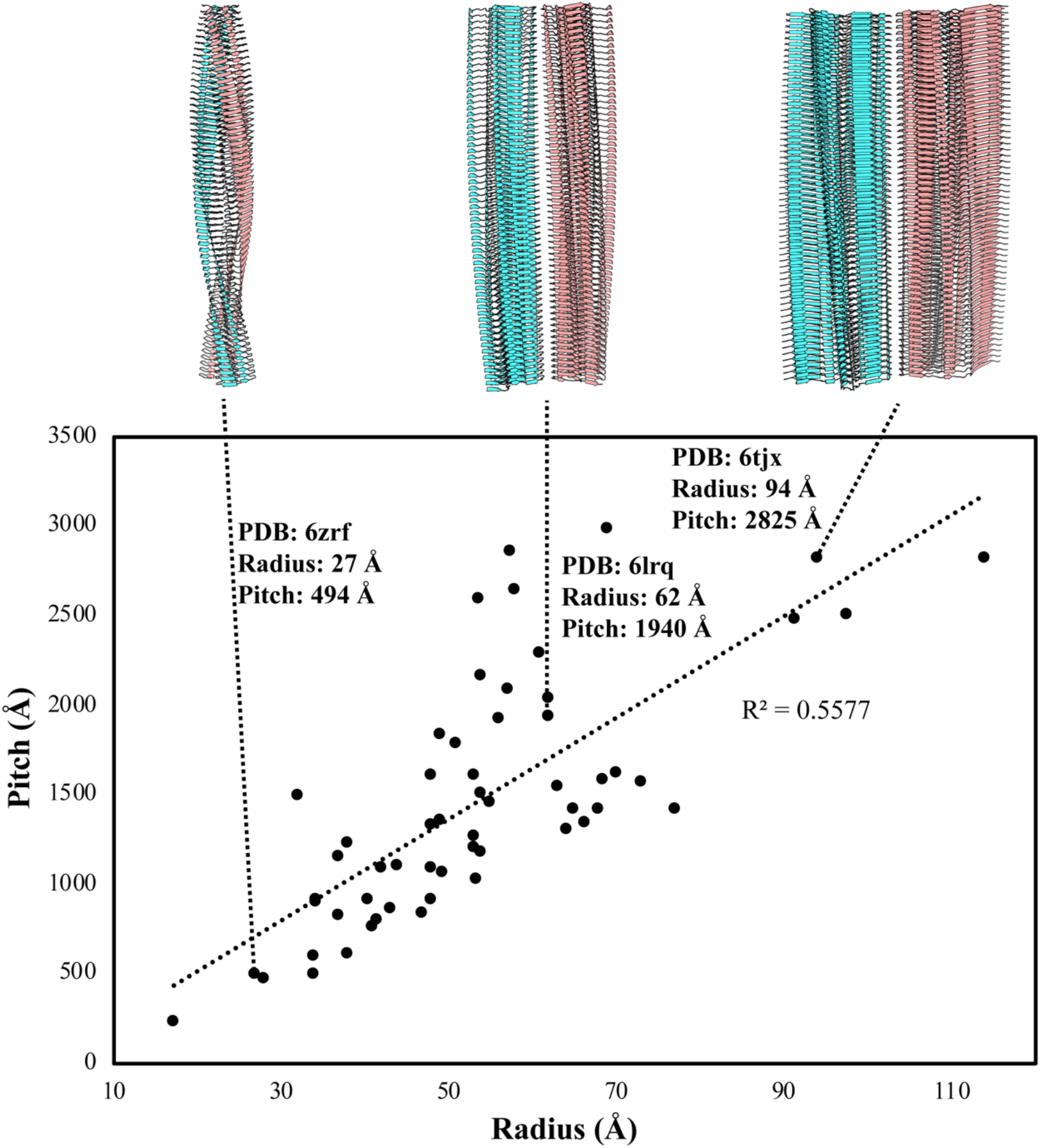
Plot of pitch versus radius for 57 cryo-EM structures demonstrates the general rule that fibrils with a larger radius tend to have a larger pitch. A trendline is shown in dashes.

### Empirical evidence that β-strand separation, hydrogen bond lengths, and off-axis tilt increase with increasing radial distance from the fibril axis

We next examined 55 fibril cryo-EM structures with atomic coordinates deposited in the PDB to test whether they support our predictions of the effect of radius on structure. In particular, we examined how fibril radius influences: i to i+1 distance for symmetrically related atoms, backbone hydrogen bonding length, and off-axis backbone hydrogen bonding tilt. We wanted to weight the reliability of our measurements by the reliability of the structure because hydrogen bond distances are sensitive to errors in atomic coordinates and cryo-EM maps of amyloid fibrils have yet to reach atomic-resolution detail. We therefore calculated and ranked all structures by their Q-score, a measure of how well the density supports each atomic coordinate (Supplementary Table 1, Supplementary Figure 1) to aid our computational analysis^12^.

Figure 6 shows example plots for four of the structures with Q-scores corresponding to an effective resolution of the model and map that is better than 2.5 Å (see Methods, Supplementary Table 1). For each structure analyzed in Figure 6 (6ufr^13^, 6xyq^14^, 6xyo^14^, and 6nwp^15^), the left most column shows the predicted distances (orange dots) and measured distances (blue dots) between symmetrically related atoms in adjacent layers in the fibril versus the distance from the helical axis for that atom pair. As Eq. 3 predicts, the distances increase at the wider parts of the fibril. The middle column measures the backbone hydrogen bond lengths in β-sheets in adjacent layers in the fibril as a function of radius from the helical axis, while the right-most column measures the tilt off the fibril axis of the same hydrogen bonds. Due to the increasing distance between symmetrically related atoms as a function of radius (left-most graph), we expect both the hydrogen bond length and tilt to increase in a manner similar to our theoretical SVQIVY helix (Figure 4). We observe that the structures in Figure 6 do indeed show modest increases in hydrogen bond length (middle plots) and off-axis tilt (right plots) as the distance from the helical axis increases. These trends support our predictions that hydrogen bonding between adjacent layers becomes weaker at regions more distant from the fibril axis, potentially placing a limit on the maximum width a fibril can grow.

**Figure 6.**
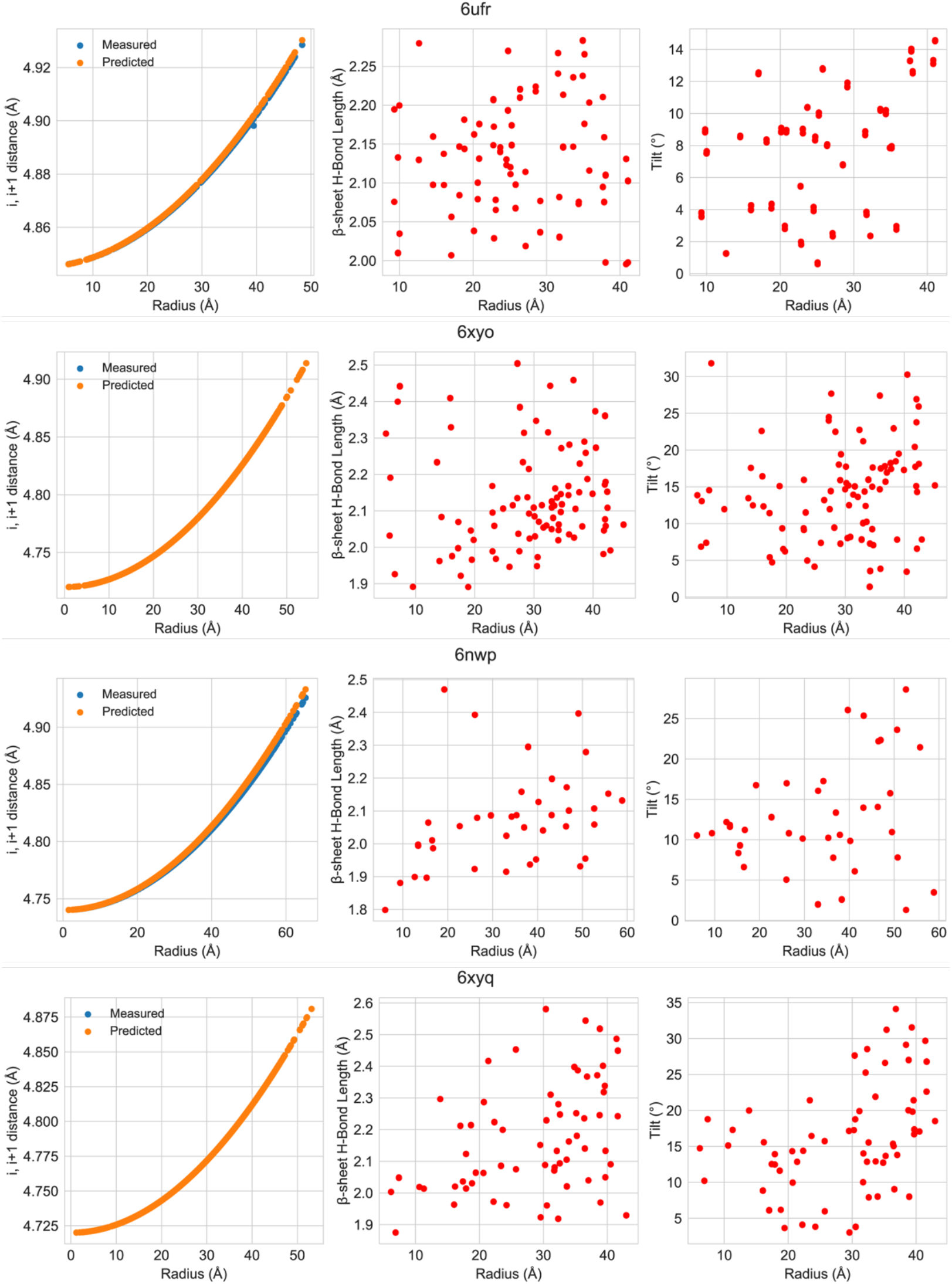
Plots of i to i+1 distance for symmetrically related atoms (left), β-sheet backbone hydrogen bond distances (middle) and off-axis tilts (right) for four of the highest resolution cryo-EM fibril structures.

Among all 55 fibril structures included in our analysis, we discovered that some trends are more strictly obeyed than others. All structures display the expected increase in distance between symmetry related atoms as the distance from the helical axis increases. However, there are some structures that show no discernible trend in the hydrogen bond length and off-axis tilt as a function of radial distance from the helical axis (Supplementary Figure 2). Figure 7 shows plots of example fibril structures with no apparent trend in the hydrogen bond length and off-axis tilt as a function of radius. Three of these structures (6zrq^16^, 6shs^17^, 6cu8^18^) have a lower predicted map/model resolution (3.4 Å, 4.0 Å, 3.5 Å, respectively) and a relatively small fibril core (34 Å, 43 Å, 48 Å, respectively). These characteristics potentially hide the trends in backbone hydrogen bond length and tilt as a function of radius because the lower resolution of the map/model potentially limits the accuracy of the atomic coordinates while the smaller fibril core limits the number of backbone hydrogen bonds that can be used in the analysis.

**Figure 7.**
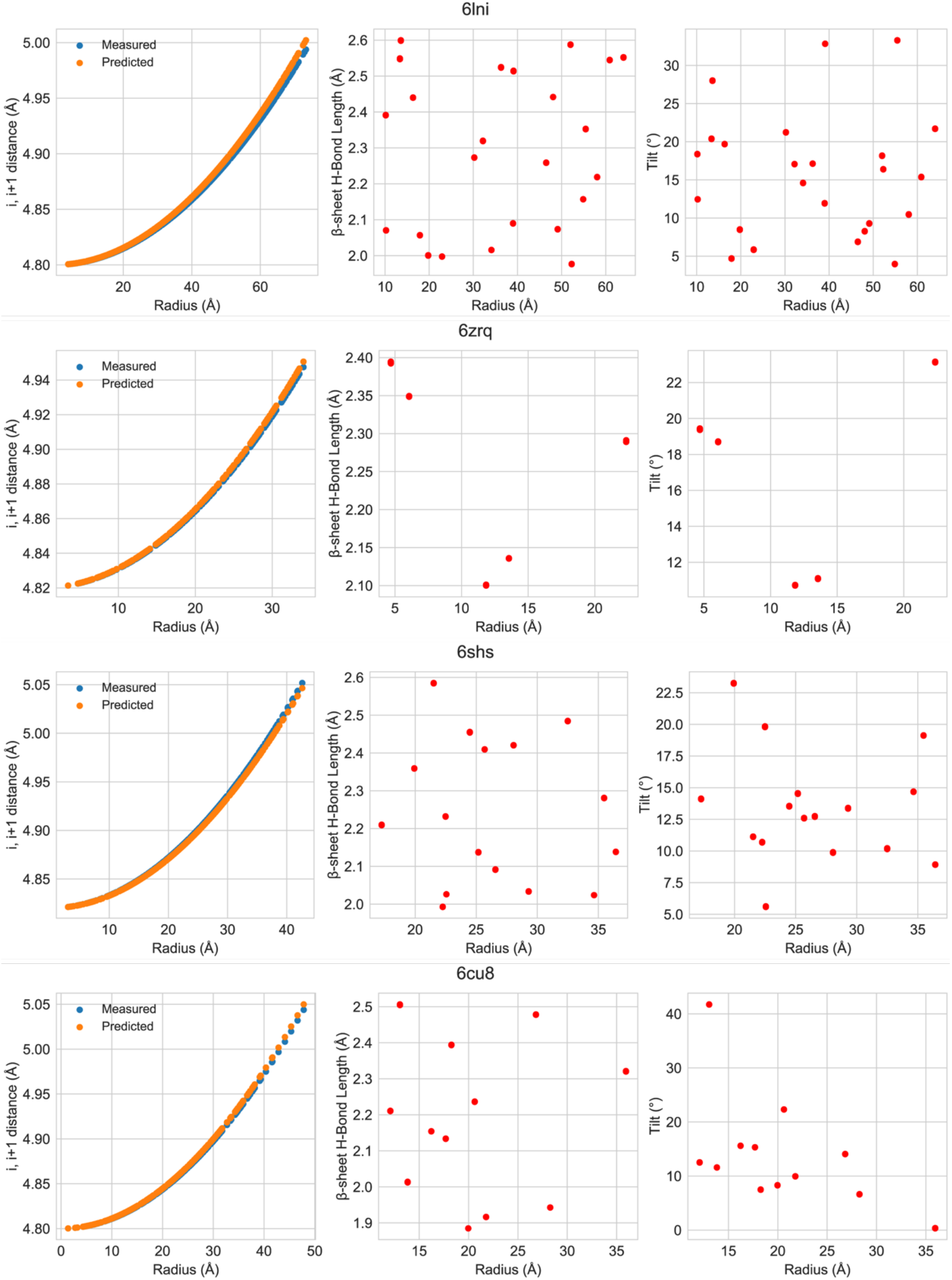
Examples of fibril structures that do not display the expected trends in β-sheet hydrogen bond length and off-axis tilt.

### Restoring Hydrogen Bond Length and Tilt Trends of a High-Resolution Fibril Structure

Unlike the previous examples with low resolution and small fibril cores, PDB 6lni^19^ has a good effective resolution (2.3 Å) and a relatively large fibril core (73 Å radius). Upon examination of the structure, it is clear that ideal restraints for hydrogen bond lengths and angles were not employed during structure refinement (Figure 8). This observation highlights the fact that even the higher resolution amyloid fibril cryo-EM maps determined to date do not have sufficient detail to accurately model all non-covalent interactions and emphasizes the need to use geometric and chemical restraints for hydrogen bonding.

**Figure 8.**
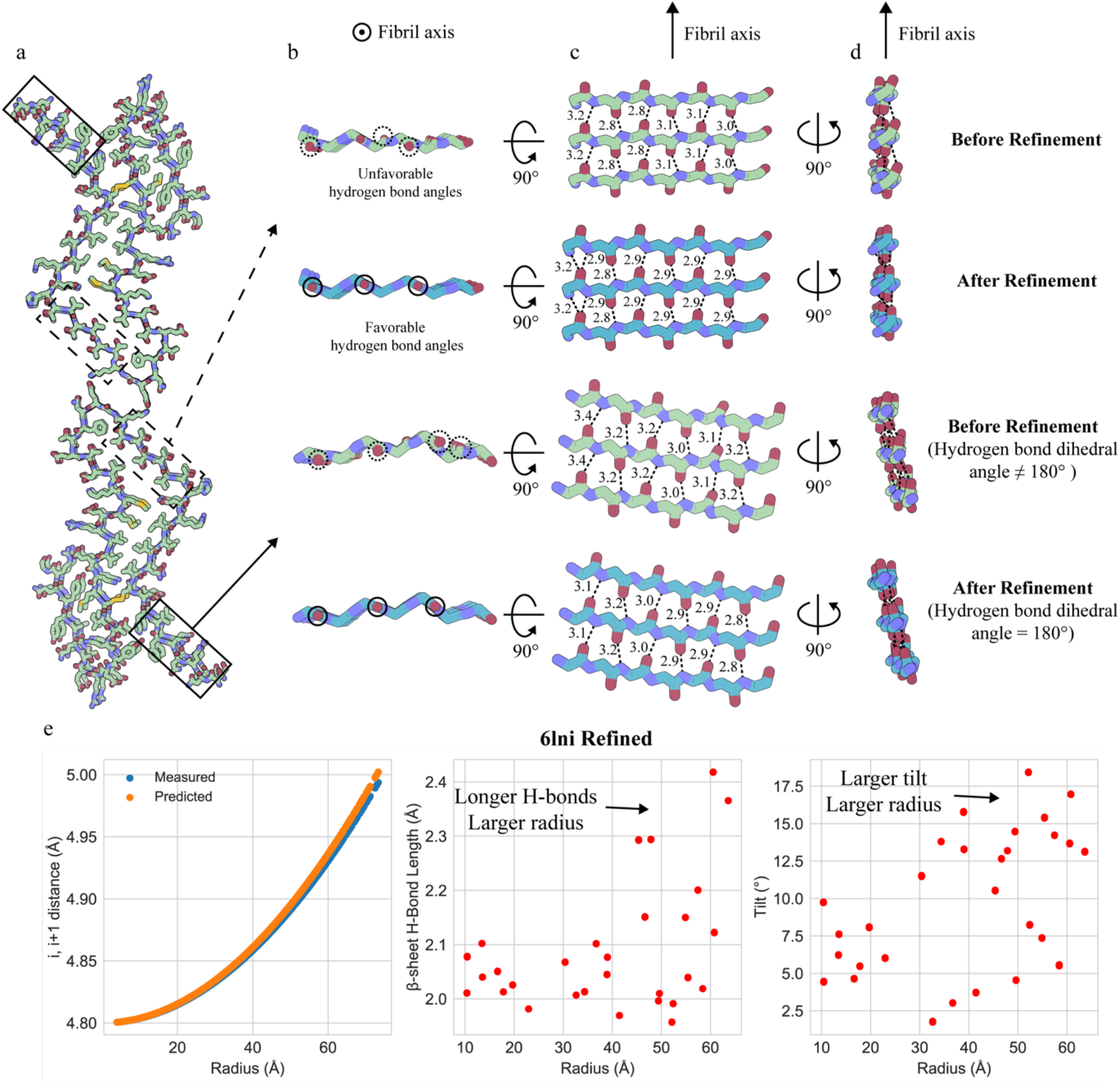
Real space refinement of PDB 6lni with hydrogen bond length and angle restraints restores predicted trends. a) Top-view of fibril 6lni formed by full-length human prion protein. b) Top-view of boxed sections showing backbone atoms before and after refinement with restraints. c) Side-view of boxed sections showing hydrogen bonding before and after refinement. d) Alternative side view of boxed sections highlighting both the correction of hydrogen bond dihedral angles and the tilting of strands that are farther from the helical axis. e) Computational analysis of interstrand distance and β-sheet hydrogen bond length and tilt as a function of radius after refinement with hydrogen bond restraints.

We tested whether β-sheet hydrogen bond distances and off-axis tilts could be restored to predicted values by refining 6lni with hydrogen bond length and planarity restraints using phenix.real_space_refine^20^. Figure 8 shows representative backbone β-sheet hydrogen bonding before and after refinement for two β-sheet regions, one close to the helical axis and one at the edge of the fibril. The refined structure shows backbone β-sheet hydrogen bonding lengths and angles agree more closely with idealized parallel, in-register β sheets (Figure 8 b-d). We then re-analyzed the backbone β-sheet hydrogen bonding lengths and off-axis tilts as a function of radius and saw that our predicted trends were more apparent (Figure 8 e) compared to the analysis of the structure from the PDB (Figure 7). This experiment highlights the importance of using hydrogen bonding restraints during structural refinement of near-atomic models to model non-covalent interactions as accurately as possible. Interestingly, the Q-score and effective resolution of the refined structure did not change noticeably from the original structure. This finding demonstrates that both geometrically correct and incorrect atomic coordinates are equally supported by near-atomic resolution cryo-EM maps and emphasizes the need for geometrical restraints on hydrogen bonding. The analysis of the refined structure supports our prediction that backbone hydrogen bond lengths and off-axis tilts increase proportionally to their distance from the helical axis of the fibril. Furthermore, it is likely that the combination of limited cryo-EM map resolution and lack of hydrogen bond restraints during refinement contributes to why some of the 55 structures in our analysis do not follow our predictions.

## Discussion

Through a combination of theory and structural analysis, we have shown that the distances between adjacent layers in amyloid fibrils increase as the fibril grows wider, and that for fibrils with a greater twist, the distances increase more steeply. These trends cause the β-sheet backbone hydrogen bonds that stabilize the fibril to grow longer, and hence weaker, towards the edges of the fibril, potentially causing fibrils to have a maximum width for a given fibril twist value. Using our analysis of hydrogen bond lengths and off-axis tilts, we identified a fibril structure with a high-resolution cryo-EM map that does not follow our predicted trends and showed that refinement with hydrogen bond length and angle restraints restored the expected hydrogen bonding patterns. This example highlights the potential utility of examining hydrogen bond lengths and off-axis tilts relative to radial distance for quality control during atomic model building into near-atomic cryo-EM maps.

Our analysis of 55 cryo-EM fibril structures indicates that the increase in hydrogen bond lengths and tilts is not as pronounced or noise-free as our theoretical SVQIVY helix (compare Figure 4d to Figure 6 and 7, Supplementary Figure 4). There are several possible reasons for this. First, the SVQIVY helix was constructed to have extra β-sheets added laterally to highlight the effects of helical symmetry on interstrand distances and hydrogen bonding. Therefore, the measurements that show the greatest effect of helical symmetry (when atoms are very far from the helical axis) will not exist in a realistic fibril since the hydrogen bonding between adjacent β-strands will be too weak (interstrand distances > ∼5.0 Å, Figure 4 c-d). Instead, only those within a range depending on the twist of the fibril will be observed in real-life fibrils, leading to the more modest trends we observe in our analysis of cryo-EM fibril structures. Second, the SVQIVY helix was based on a high-resolution microED crystal structure where the position of the atoms involved in hydrogen bonds are known to a high accuracy. This is unlike cryo-EM fibril structures that have lower resolution, making the positions of atoms approximate and necessitating great care in modeling hydrogen bonds. Third, in crystal structures of short segments of amyloid proteins, deviations from canonical β-strand conformation are rare and sheets are flat, so the backbone geometry is rather uniform. Most amyloid proteins are much longer and exhibit greater diversity of residue conformation with varying degrees of β-strand twist, which make our backbone hydrogen bond measurements noisier in comparison to a theoretical helix based on a high resolution crystal structure.

In addition to the strain the helical twist imposes on the perimeter of fibrils, there may be other factors that limit the width of amyloid fibrils. One such factor is that not all amino acids are equally amyloidogenic. In other words, some sequences tend to not to be compatible with the cross-β fold^21^. For instance, proline residues are incapable of participating in the backbone hydrogen bonding of β-sheets, and excessive residues of unbalanced charge would incur electrostatic repulsion in the cross-β fold due to the 4.8 Å stacking of identical amino acids in parallel, in-register β-sheets along the fibril axis.

A good example for this situation is the protein tau, which forms amyloid fibrils in Alzheimer’s and other diseases^15, 22–26^. The longest isoform of tau is 441 residues, with a large N-terminal sequence that is highly positively charged and proline rich, four C-terminal microtubule binding pseudo-repeat domains containing known amyloidogenic sequences, and an additional ∼40 residue far C-terminal domain. In the fibril structures of tau, 60-90 residues of the repeat domains form the fibril core, while the remaining 300+ residues from the N- and far C-terminus are disordered in the fuzzy coat. This is consistent with the fact that the N-terminus is proline-rich and highly charged, but interestingly, the amyloid-segment predicting software ZipperDB^21^ indicates there are amyloidogenic segments in the ∼40 residues of the far C-terminus. However, in all the tau fibril structures determined thus far, these additional amyloidogenic sequences do not contribute to the fibril core.

We demonstrated that there is a correlation between fibril twist and width; namely, that fibrils with a large twist must have smaller cores, while those with a smaller twist can have larger cores (Figure 5). However, it is not clear whether it is the twist that informs the size of the core, or if the size of the core determines the helical twist. Chou and Scheraga used computational chemistry techniques to show that β-strands favor a right-handed twist (more so than left-handed or flat) due to the steric interactions of i to i+2 side chains in L-amino acids; however, the twist of the β-strands is reduced in β-sheets – compared to isolated β-strands – due to interchain interactions (e.g., backbone hydrogen bonding, side chain-side chain interactions) “flattening” the twist of the constituent strands^8^. The large number of β-sheet residues in an amyloid fibril core therefore leads to many interchain interactions (e.g., backbone hydrogen bonding) that serve to flatten the twist of the constituent β-strands leading to extremely small twists (∼0.5° – 3.5°, Supplementary Table 2) compared to the large twists (>10°) seen in β-sheets in globular proteins. Therefore, we hypothesize that it is the number of residues that participate in forming the fibril core that determines the twist of the amyloid fibril. This leads to smaller cores having larger twists and larger cores having smaller twists. Then for a given fibril core size, the twist of the fibril will destabilize the β-sheets at the edges of the fibril, leading to either i) less residues from the constituent protein chains being able to be added or ii) the inability of additional protofilaments to be added laterally to the fibril.

For scenario (i) above, one could imagine a theoretical protein sequence of 1000 residues where every residue could adopt the cross-β fold. Given the conclusion of Chou and Scheraga, if all 1000 residues of our imaginary protein formed part of the fibril core, the twist of the fibril could approach zero since the amount of interchain interactions favoring flattened β-strands would outweigh the i to i+2 side chain steric interactions that favor a twisted β-strand. Further study would be needed to see if this were indeed possible, or if a strain in the polypeptide chain exists that will cause it to twist, leading to a limited width for the fibril.

A potentially illuminating example for the above situation is the fused in sarcoma (FUS) protein. FUS is a 526 residue protein with a 214 residue N-terminal low complexity domain (LCD). Recent studies have shown that, when produced separately, both the N- (2-108) and C-terminal (111-214) halves of the LCD can form fibrils, while the full LCD forms fibrils with only the N-terminal core^27, 28^. This raises the question why a larger core, containing both the N- and C-terminal LCD, does not form. The authors in Lee, et al. correctly point out that if a putative fibril formed having both the N- and C-terminal LCD fibril cores present, and the fibril followed the same helical twist as the C-terminal-only fibril, the interstrand distances in the N-terminal core (the periphery of the fibril) would become too great^28^. However, it is possible that in a fibril where both segments contribute to the core, the helical twist may become smaller due to the additional interstrand interactions flattening the strands, thereby permitting a wider fibril. Or a fibril may form with a new fibril core structure where both N- and C-terminal LCD segments interact and the twist similarly becomes smaller since there are more interchain interactions flattening the fibril. The fact that neither of these scenarios occur, at least under tested experimental conditions, may be evidence that fibrils cannot grow indefinitely wide since there may be some amount of twist necessary to prevent straining the polypeptide chains, hence limiting the width of the fibril. This may be similar to tau where even in the largest diameter tau fibril^26^ (PDB 6tjx; diameter 388 Å), with a twist of -0.61°, containing all four microtubule repeat domains (residues 274-380) there exist residues (381-441) just beyond the C-terminal end of the core that are predicted to be amyloidogenic by ZipperDB that are nonetheless not included in the fibril core.

Numerous studies have identified apparently non-twisting amyloid fibrils. For example, Schweighauser, et al. and Li, et al, identified non-twisting fibrils of alpha-synuclein from Parkinson’s Disease and from recombinantly assembled wild-type fibrils, respectively^14, 18^. Due to their lack of helicity, their structures are unable to be determined by current cryo-EM helical reconstruction methods (although ssNMR would be ideal for structure determination since it does not rely on the helicity of fibrils). At present, it is unknown what factors lead these fibrils to not twist. Liberta, et al. demonstrated right- and left-handed serum amyloid A protein amyloid fibrils from human and mouse, respectively^29^. Their analysis of Ramachandran angles showed that right-handed fibrils have a slight majority of residues in left-handed β-strand conformation and vice versa for the left-handed fibrils. Therefore, one possible explanation for non-twisting fibrils is that there is an equal ratio of right- and left-handed β-strands in the protein chains, although this has not yet been shown experimentally.

Our theory predicts that the helicity of amyloid fibrils constrains their width; however, this leads to the question: why do the observed non-twisting fibrils not associate laterally (by translational, rather than helical symmetry) to create ordered two- or three dimensional (crystalline) arrays? One possible explanation for this is that the external surfaces of fibrils are not amenable to cross-β association. This could be due to the fact that charged residues tend to prefer to interact with water and remain solvent exposed rather than be buried inside the fibril core, as in globular proteins. However, there are numerous cases, including recombinantly assembled wild-type^30^, E46K^13^, and Tyr39p^31^ alpha-synuclein, and full-length human prion protein^19^ fibrils where protofilaments assemble strictly through electrostatic interactions. This suggests that fibrils can indeed extend their width through lateral association of protofilaments mediated by electrostatic interactions, and that in these cases, the helical twist may impose the limit on width and not the surface characteristics of the fibril. Given these examples, it is unclear why non-twisting fibrils have not yet been observed to associate laterally. Perhaps due to their rarity, a fibril surface composition for non-twisting fibrils has not yet been sampled that may allow more extensive lateral growth. Further work determining the structures of non-twisting fibrils may illuminate the specific reasons these fibrils do not grow wider. In conclusion, our analysis suggests that fibril width is controlled by a multitude of factors, including helicity, the amyloidogenicity of the protein sequence, and the outward facing surfaces of the fibril.

Understanding the rules that determine the width of fibrils may enable the design of fibrils of arbitrary width. Here, we have shown that the twist of the fibril plays an important role in controlling the fibril width; thus, methods to control the twist may present a viable path to controlling fibril width. For instance, the use of alternating L- and D-amino acids may lead to fibril backbones that do not twist, therefore forming non-twisting fibrils that could have a larger width. It has already been shown that fibrils have comparable mechanical properties to that of steel and silk^32, 33^. The addition of more amino acids to a given fibril core would add to the network of interactions that stabilize fibrils and could potentially increase the mechanical stability of amyloid fibrils. Furthermore, fine-tuning the width of a fibril may affect its assembly/disassembly kinetics which may be useful for drug delivery methods that rely on controlled release of monomers from fibrils^34^. At present, it is not possible to rationally encode a fibril structure from a designed amino acid sequence, although meta-analyses such as the one we conducted here may hold clues to learning the design principles of amyloid fibrils, possibly enabling structure prediction and/or structure design as has been accomplished with globular proteins^35^.

## Author Contributions

D.R.B conceived of and oversaw the project. D.R.B., N.A.M., and M.R.S wrote the software for computational analysis. D.R.B. wrote the paper with input from all authors.

## Acknowledgements

D.R.B was supported by the National Science Foundation Graduate Research Fellowship Program and the UCLA Graduate Division Dissertation Year Fellowship.

## Material and Methods

### Preparation of pdb and map files

All pdb and map files were downloaded from the Protein Data Bank (See Supplementary Table 2 for full list). In order to allow for easy computation, we curated the files as follows. We moved all fibrils so that their helical axis was coincident with the z=0 axis. We also removed all alternate rotamers. For many fibrils, we observed that the pdb file did not follow the exact symmetry of the map file. This is likely due to the fact that refinement programs such as phenix.real_space_refine, although they do use non-crystallographic symmetry, do not impose helical symmetry on the refined pdb. Therefore, we used the helical twist and rise values listed in the corresponding publication for each structure and helicized the model before our analysis using pdbset^36^.

For the analysis of the pitch and radius of the Pick’s Disease Wide Fibril used in Figure 5, we downloaded the Wide Fibril map and Narrow Fibril model from the PDB and built a hypothetical Wide Fibril model as the authors propose in Falcon, et al. by rigid-body fitting two Narrow Fibrils into the Wide Fibril map^23^. We measured the maximum radius of the theoretical Wide Fibril and calculated its pitch using the Wide Fibril helical parameters from Falcon, et al. For the RipK3 fibril^11^, we included radius and twist parameters in Fig. 5, but did not calculate hydrogen bonding parameters due to the low resolution of the cryo-EM map and the resulting uncertainty in the atomic model.

### Q-score analysis

We measured the Q-score for all maps and models after moving the fibril axis to the origin and helicizing all models according to their published helical parameters. We converted Q-scores into effective resolution by the equation Q-score = -0.178 * (Expected Resolution)+1.119 from ref.^12^ (Supplementary Table 1). We plotted the reported resolution from the publication versus the effective resolution from the Q-score in Supplementary Figure 1. This indicated that most structures have a better effective resolution than reported in the publication, possibly suggesting the gold-standard FSC slightly underestimates the resolution for amyloid fibril reconstructions or that the Q-score analysis needs to be re-calibrated for amyloid fibril structures compared to globular proteins.

### Computational analysis

We wrote a FORTRAN program that measures the maximum distance from the helical axis for every atom in a fibril in order to create Figure 5. Pitch values for Figure 5 were calculated from the twist and rise values in the publications. In order to calculate the i to i+1 distance between symmetrically related atoms, we wrote a Python program that identified i to i+1 chains (symmetrically related chains immediately above or below one another in the helix) and calculated the distance between identical atoms in the two chains, as well as the distance from the helical axis for those atoms. In order to calculate the backbone-backbone hydrogen bonding distance, we added hydrogens to all fibril structures using phenix.reduce^20^. We then wrote a Python 3 program to identify all backbone-backbone hydrogen bonds between amide hydrogens and carbonyl hydrogens with distance greater than 1.6 Å and less than 2.6 Å. We also measured the off-axis tilt for each hydrogen bond by measuring the magnitude of the carbonyl C-O bond vector projected onto the X-Y plane. The arccos of this value divided by the magnitude of the C-O bond vector in all three dimensions, gives the complementary angle to the bond tilt relative to the fibril axis. We further calculated the Ramachandran angles of all residues in the fibrils so that we could explicitly examine β-sheet hydrogen bonding – where both hydrogen bond donor and acceptor are in β-strand conformations.

### Code availability

All code used for calculating fibril radii, interstrand distances, hydrogen bond distances and tilts, centering cryo-EM maps and models, generating hydrogen bond distance and angle restraints for real space refinement, and helicizing fibril models are available on the following GitHub repository https://github.com/nikashmyn/Amyloid_fibril_width.git.

## Supplementary Information

**Supplementary Table 1.**
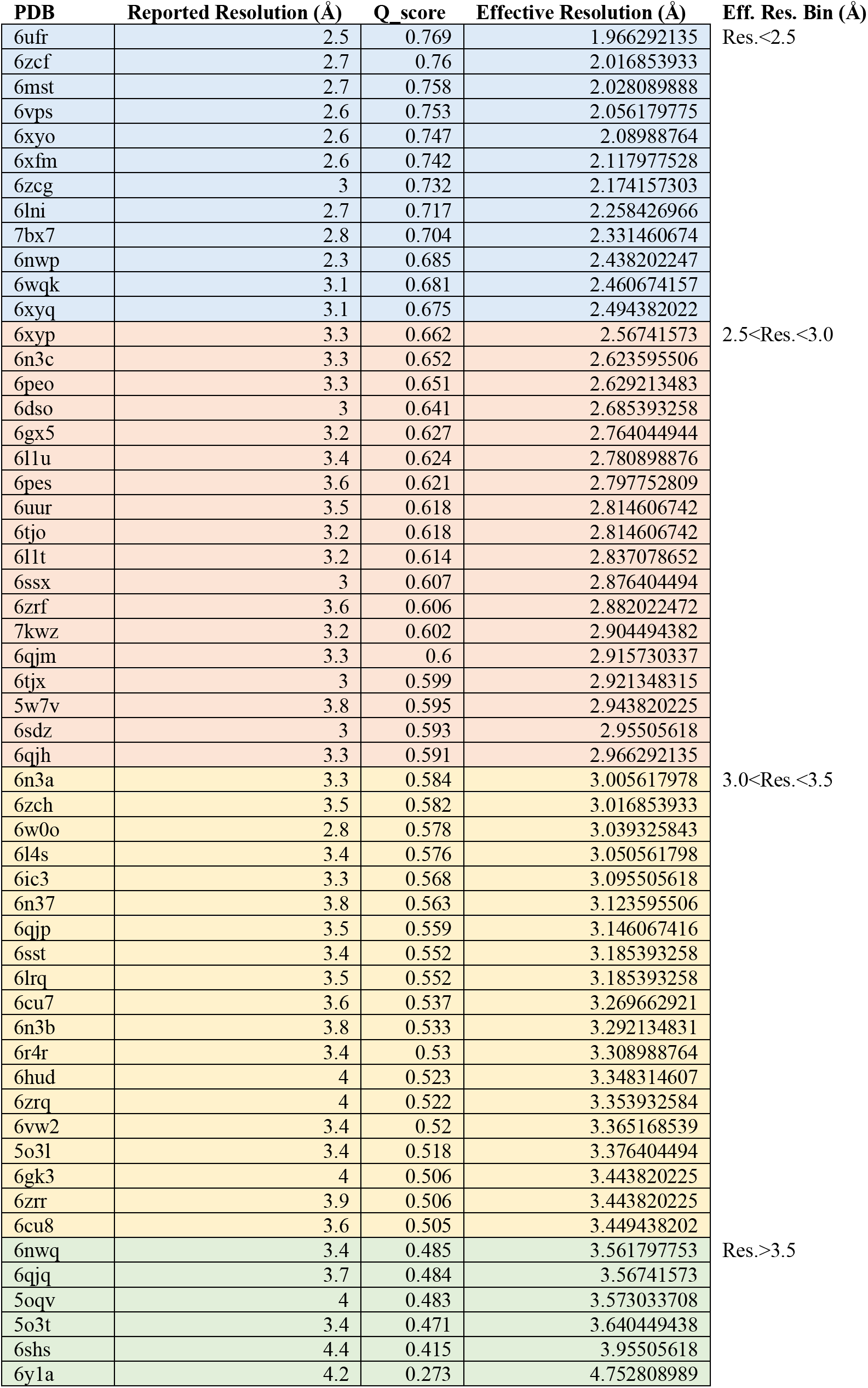
Q-score analysis of all 55 cryo-EM structures

**Supplementary Figure 1.**
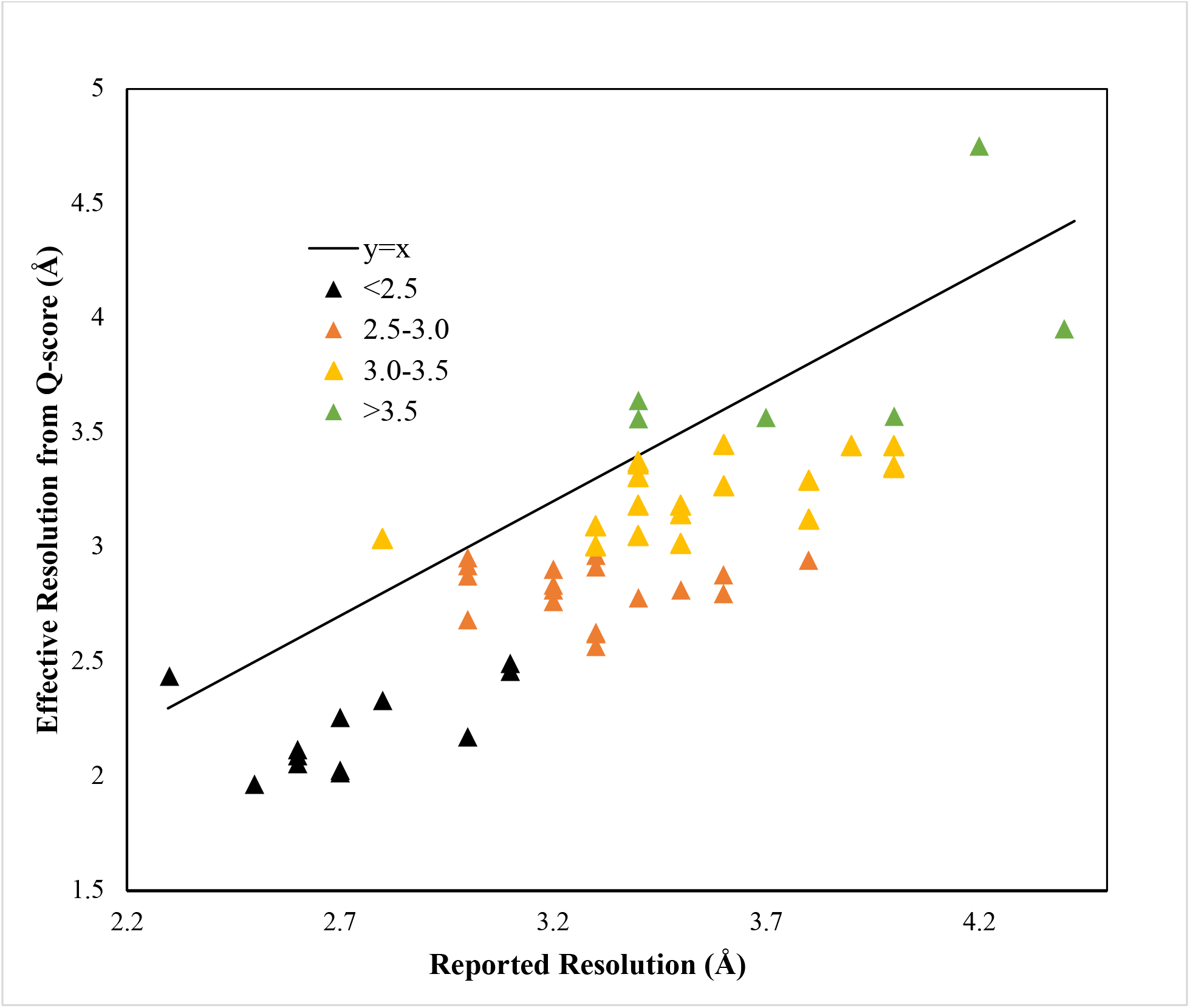
Plot of reported resolution from publication versus expected resolution from Q-score analysis (Q-score = -0.178 * (Expected Resolution)+1.119).

**Supplementary Figure 2.**
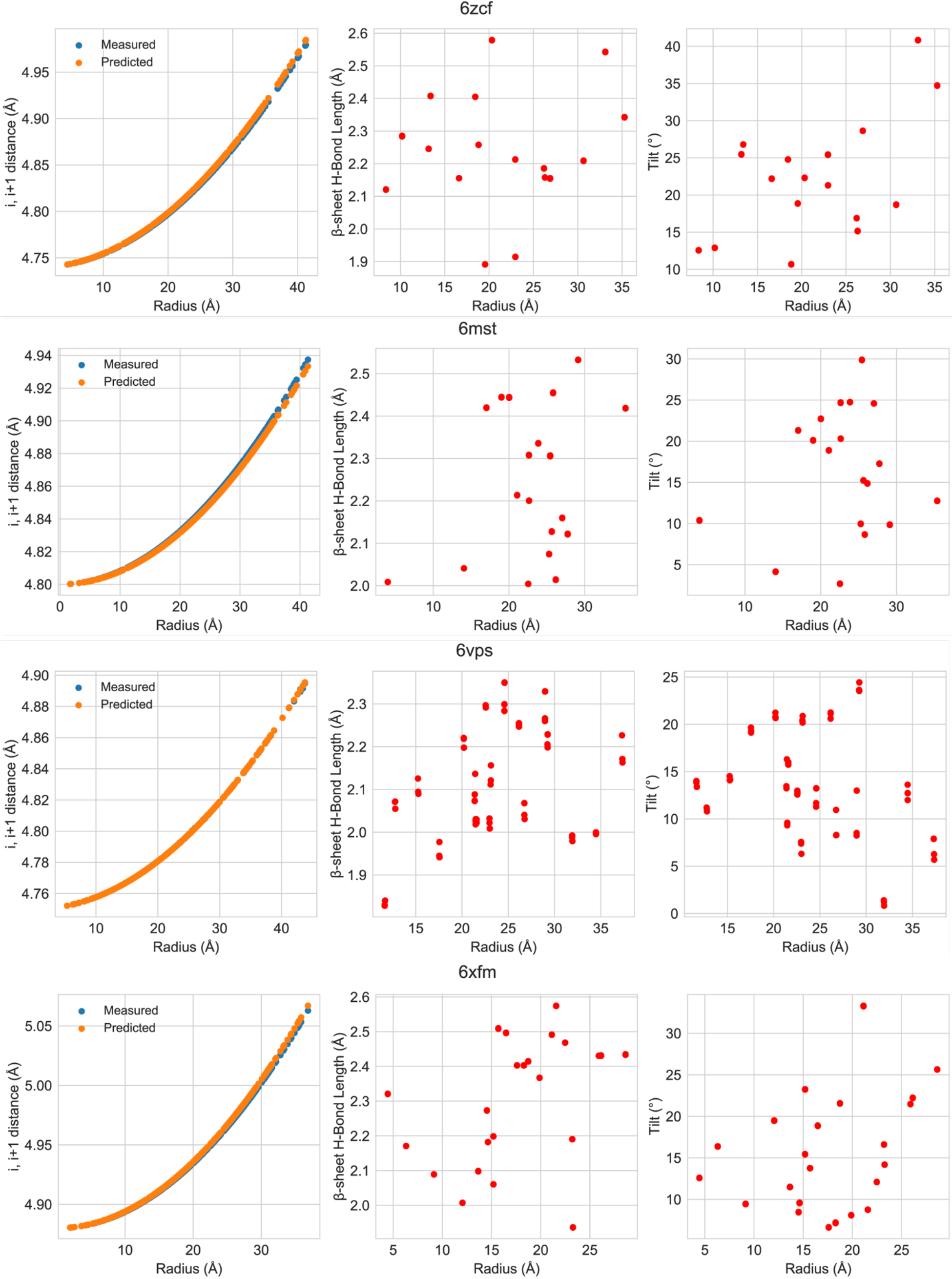
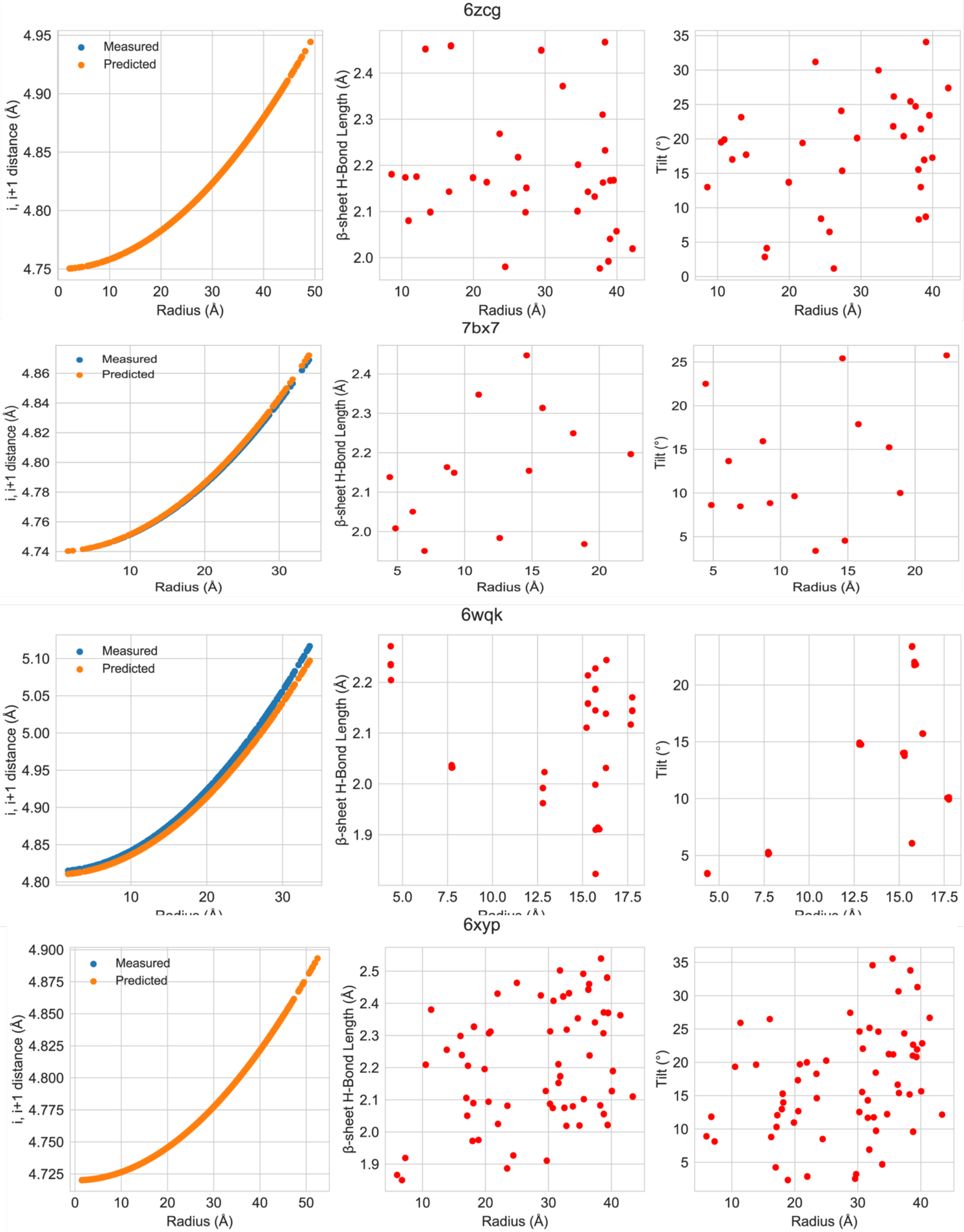
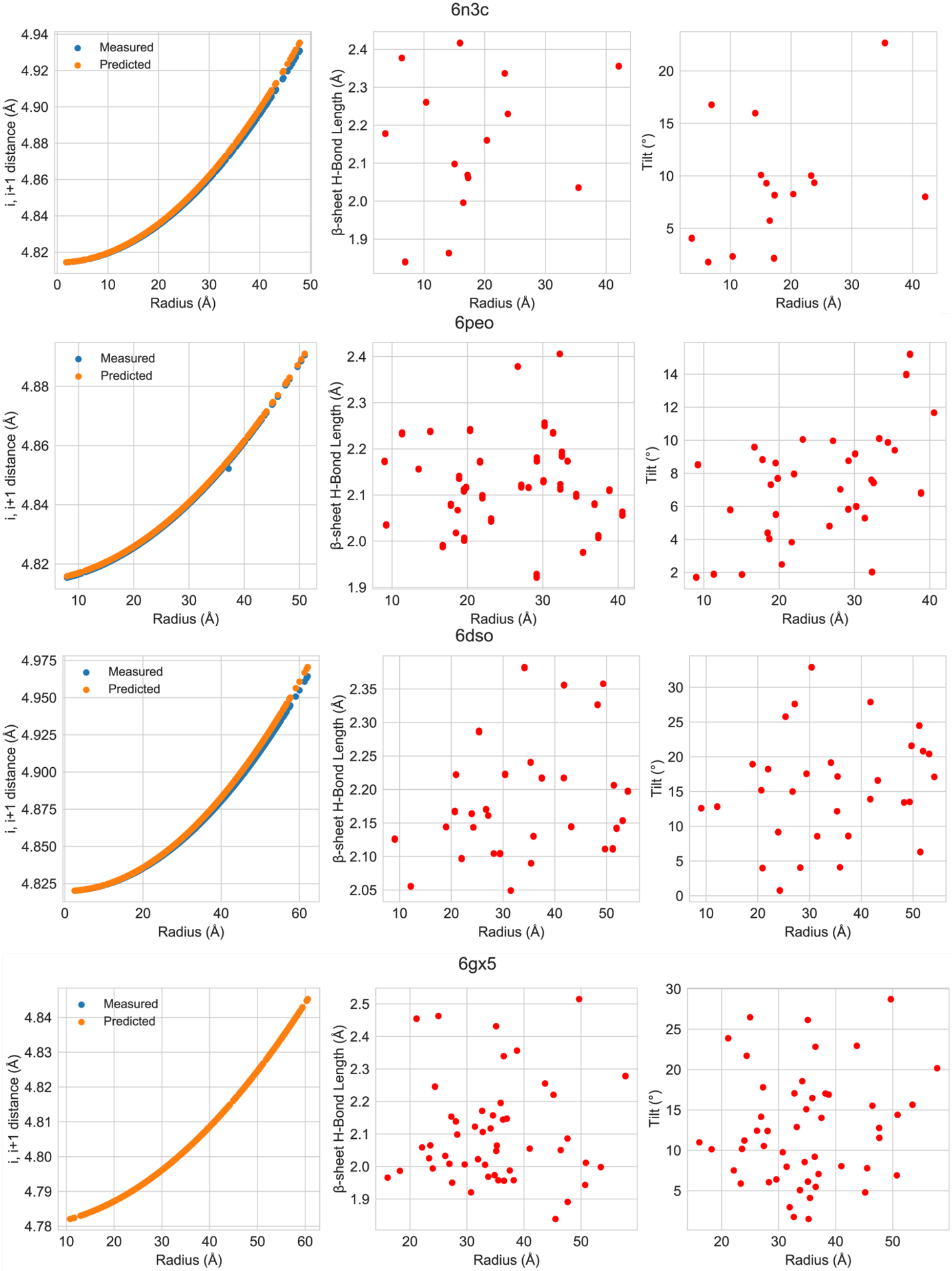
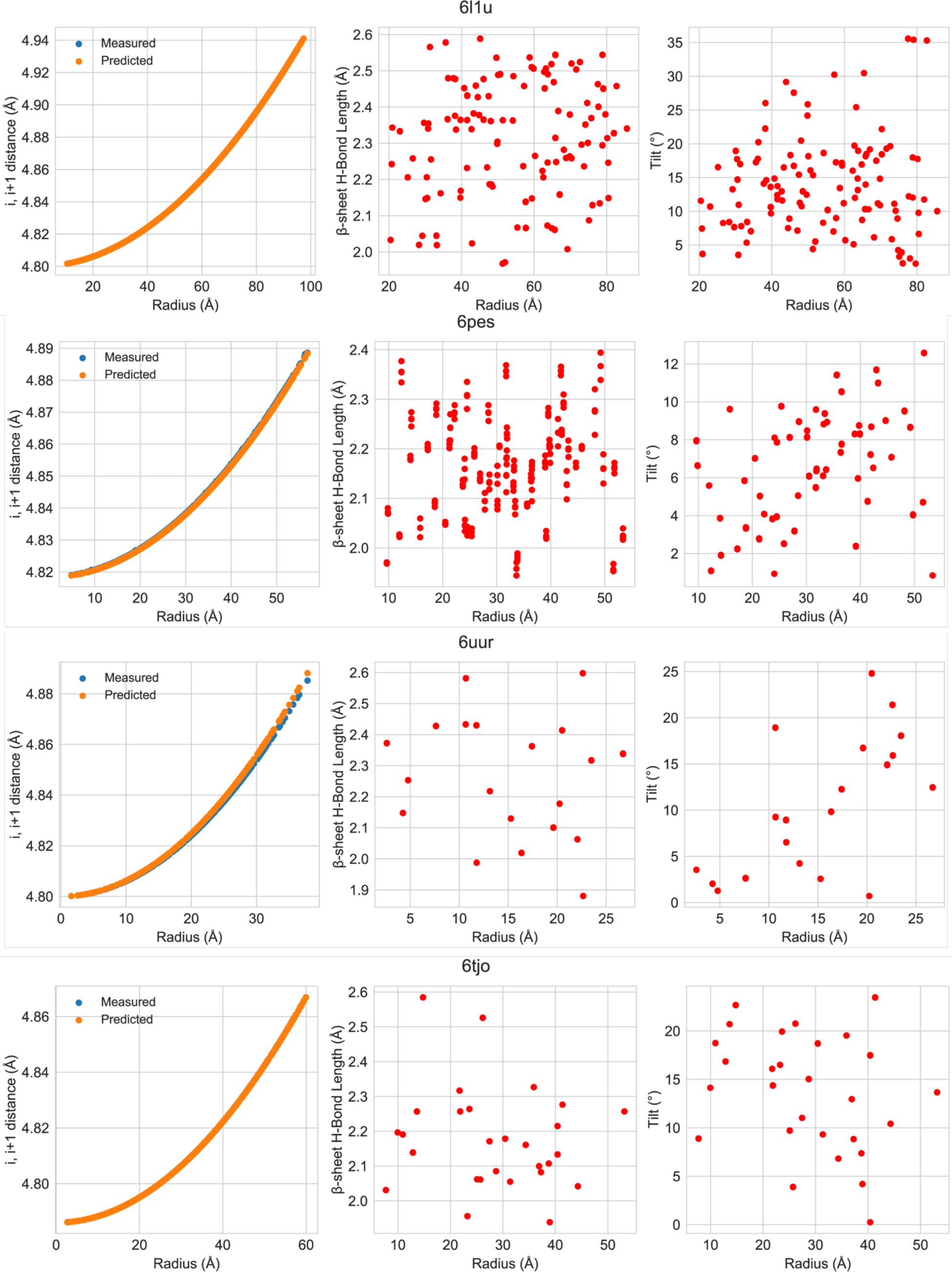
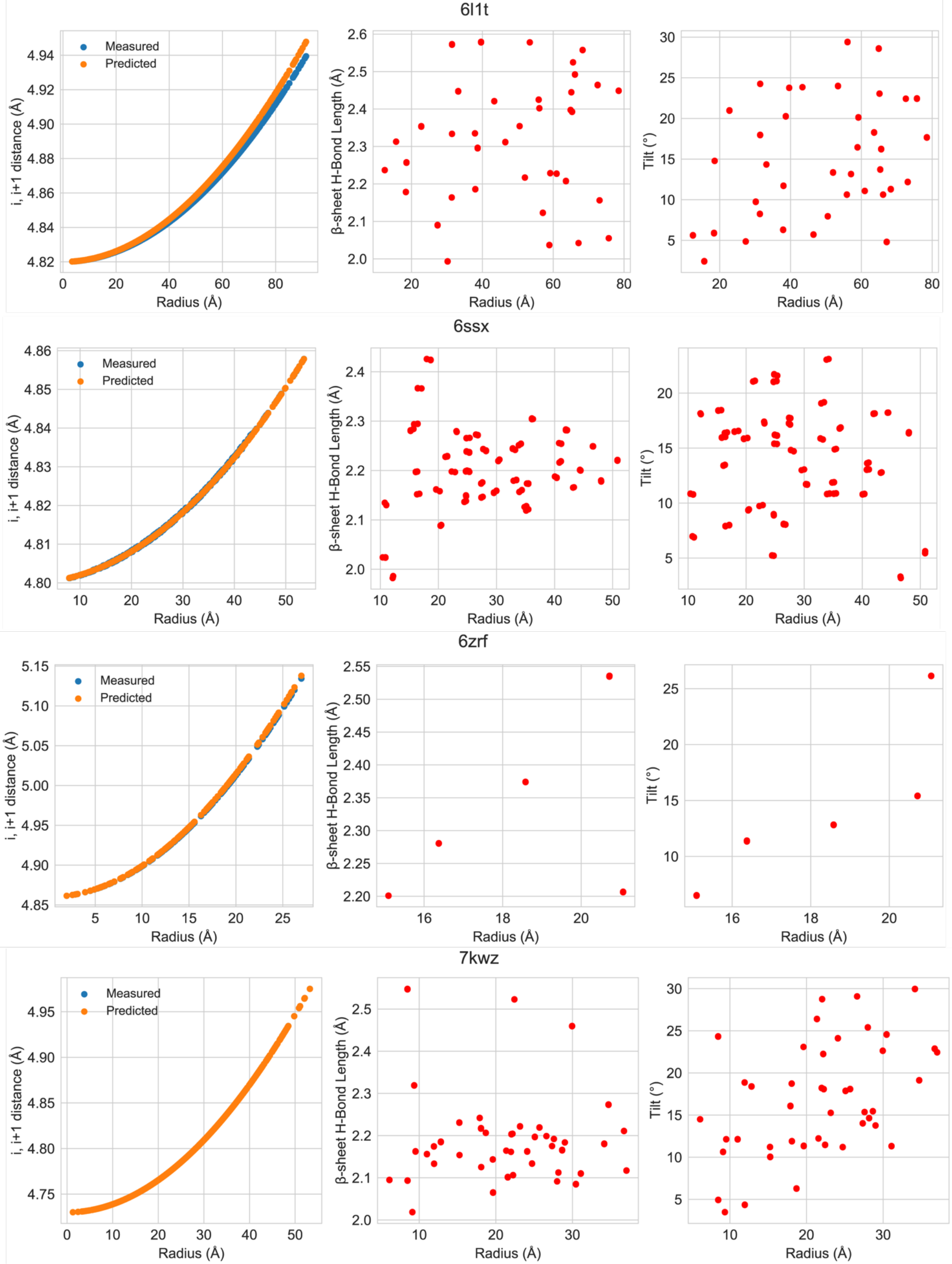
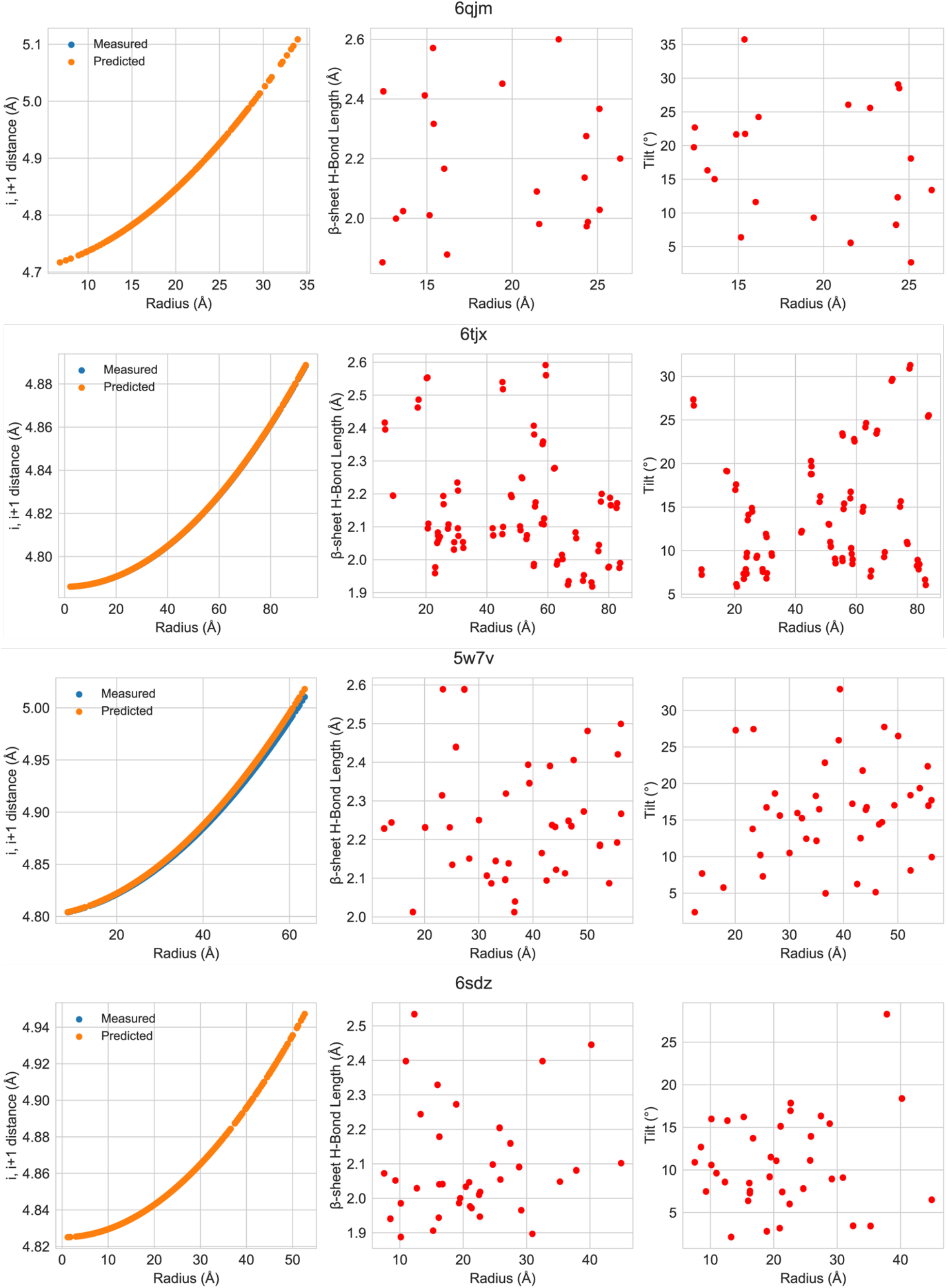
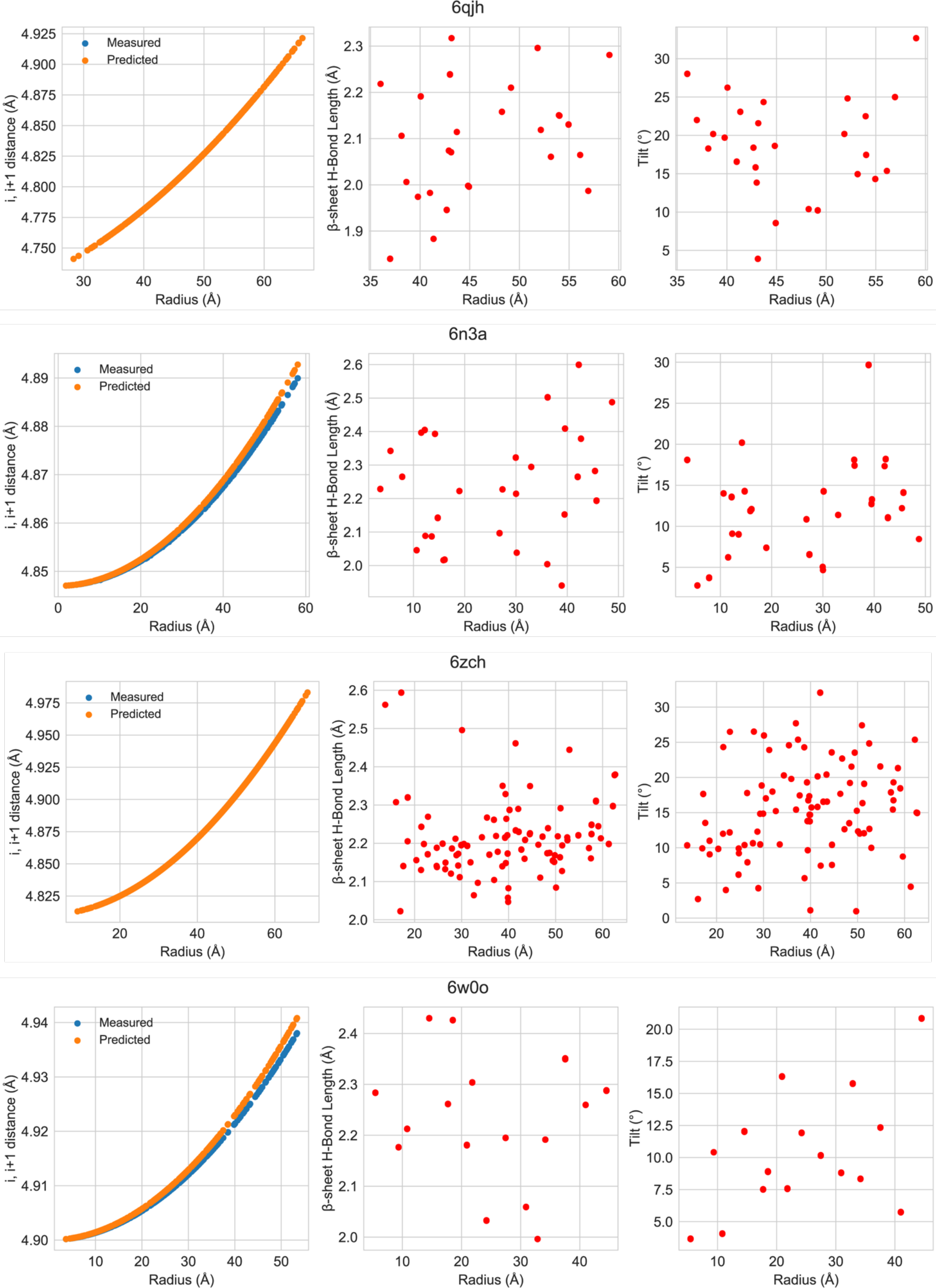
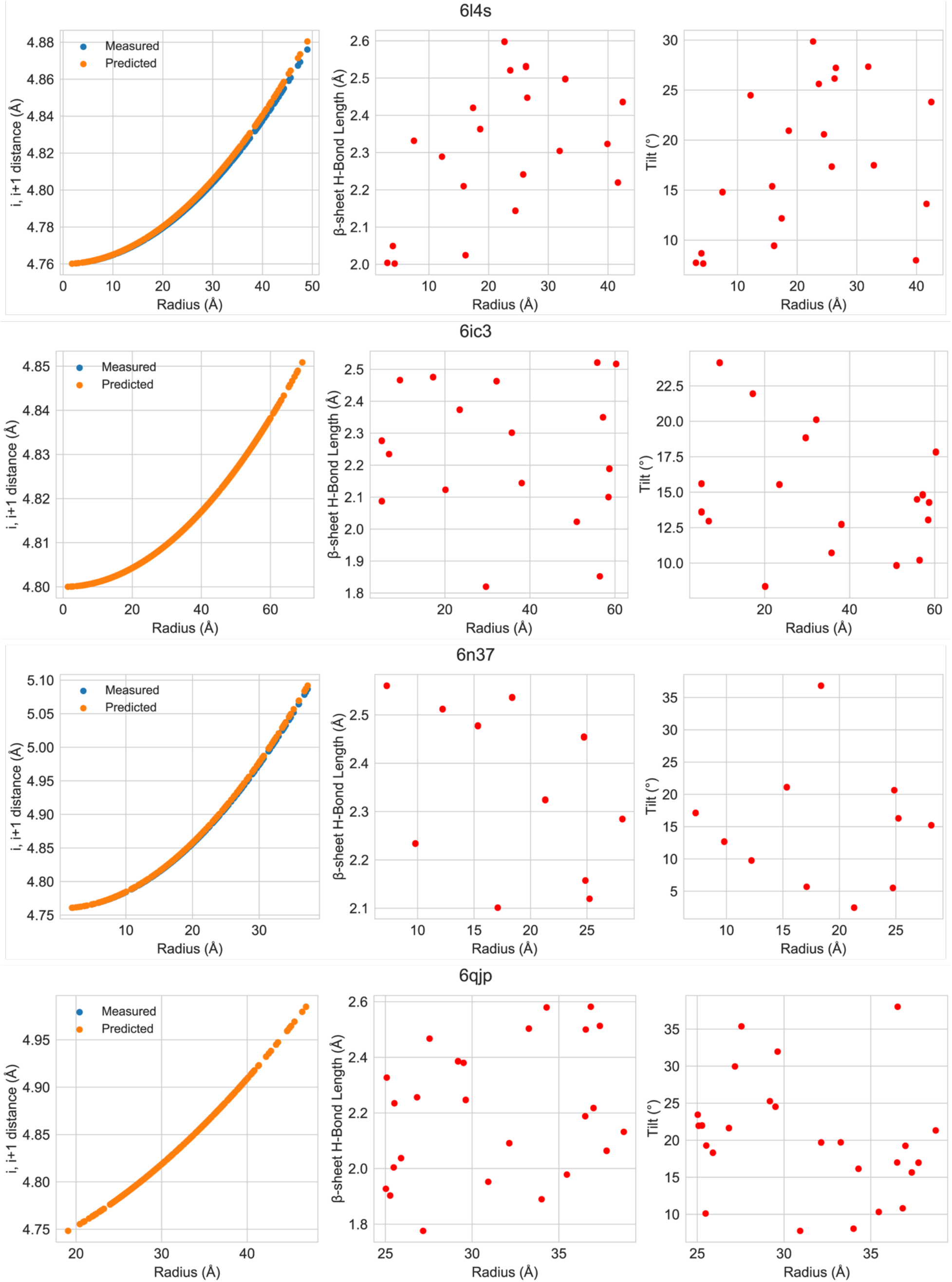
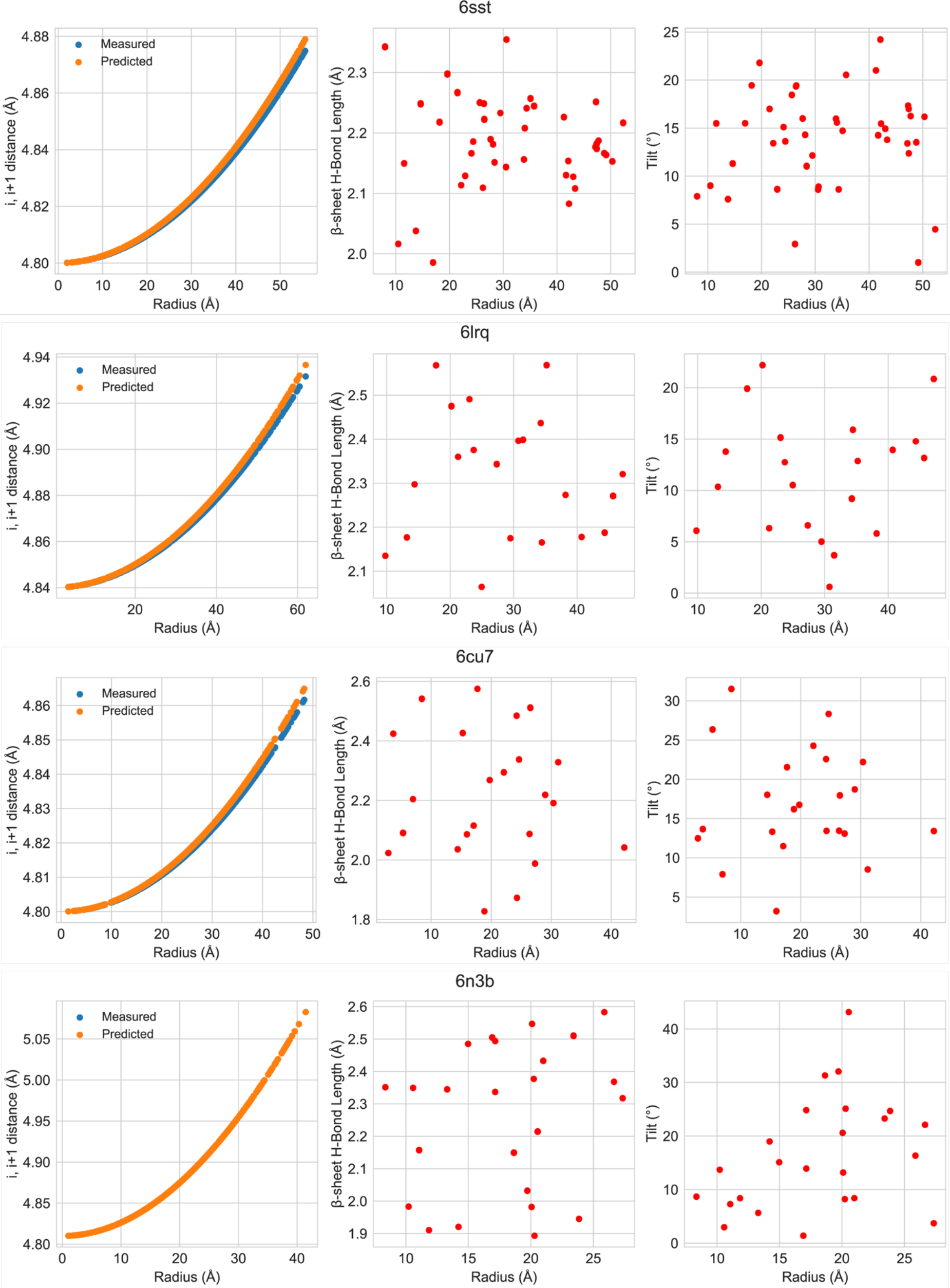
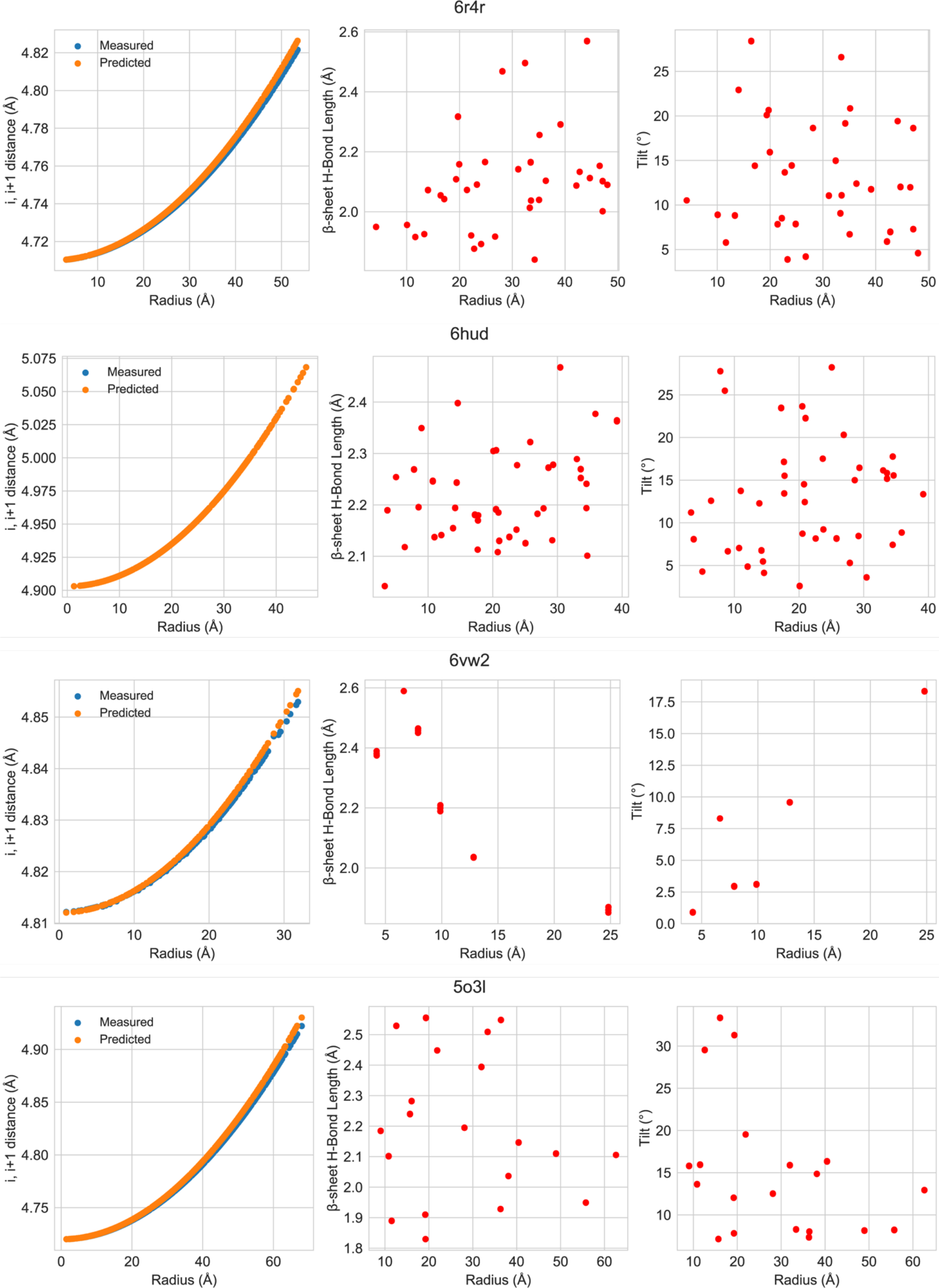
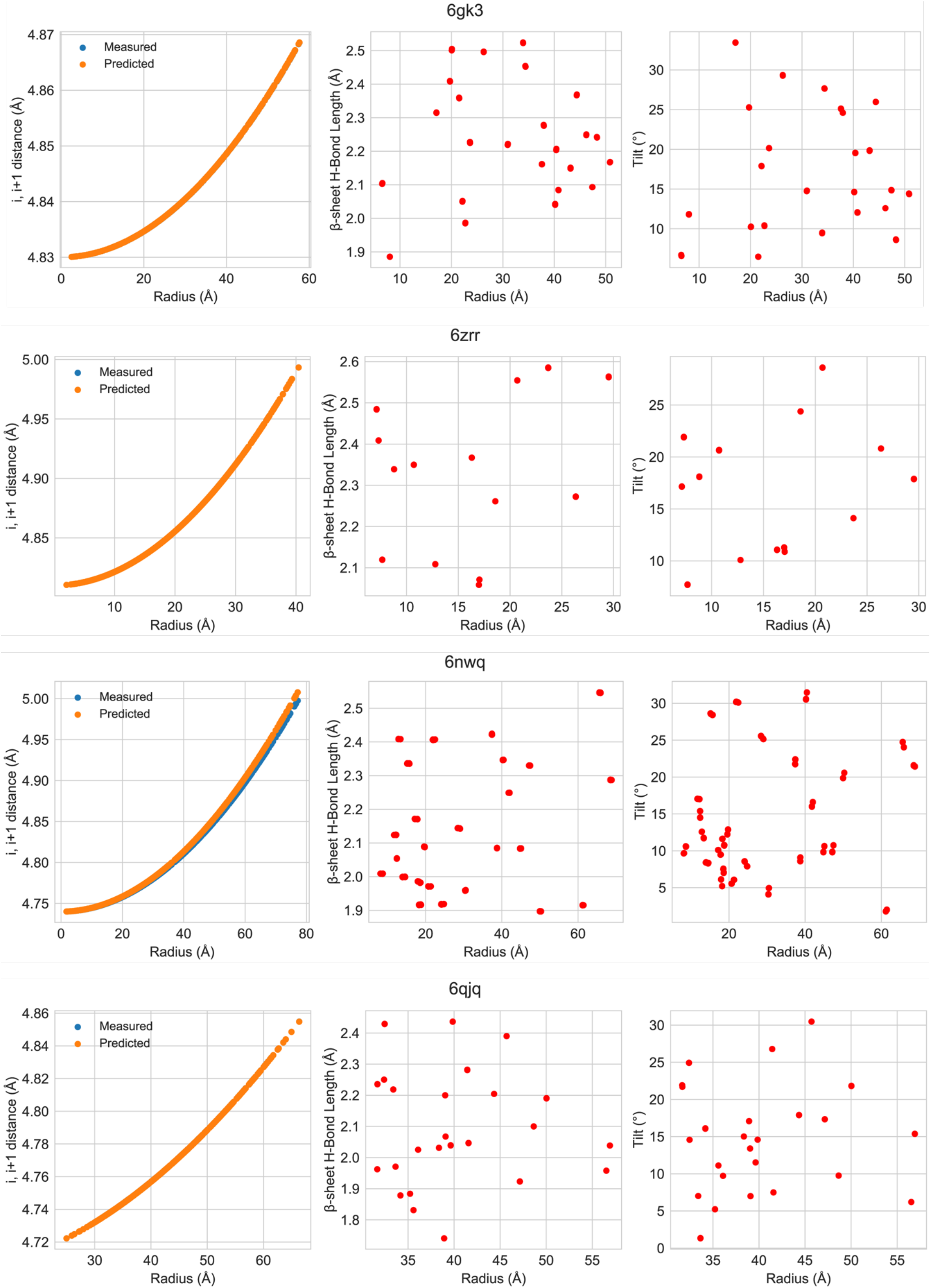
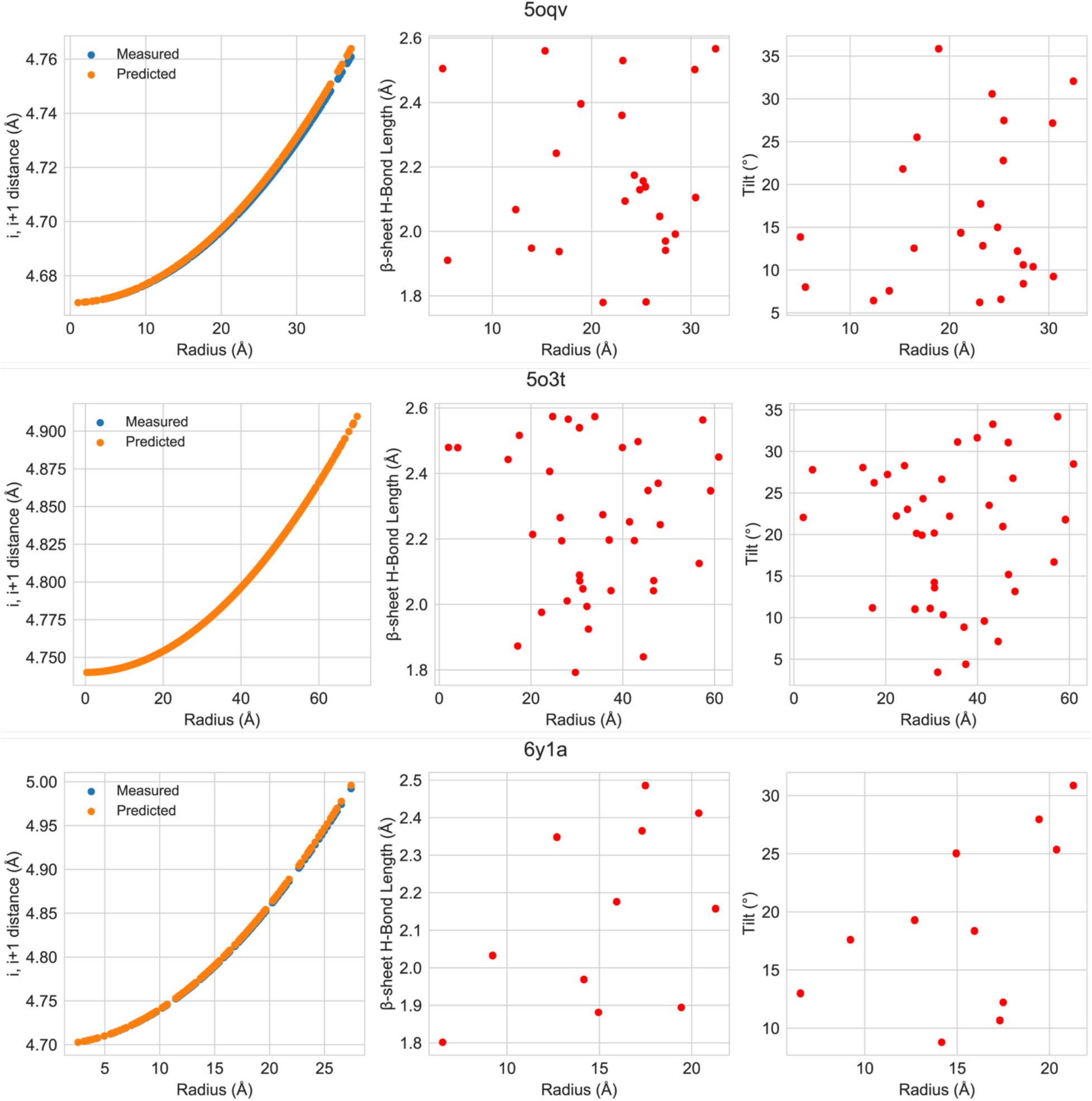
Plots of fibril structures showing i, i+1 distance between symmetry related atoms, backbone hydrogen bond distances and off-axis tilts between i, i+1 β-strands.

**Supplementary Table 2.**
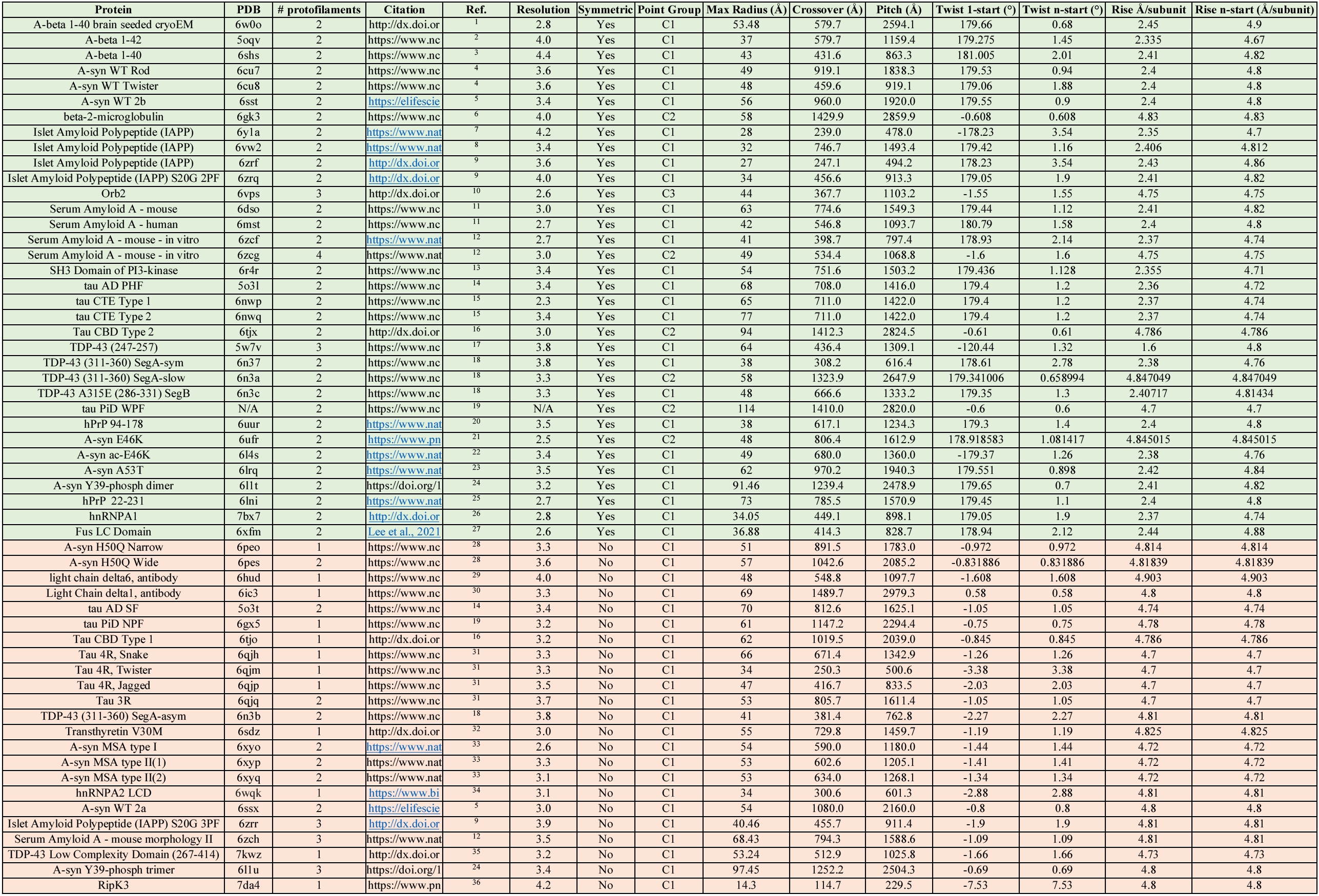
Information for all structures used in analysis.

